# Isolation, Development, and Genomic Analysis of *Bacillus megaterium* SR7 for Growth and Metabolite Production Under Supercritical Carbon Dioxide

**DOI:** 10.1101/352807

**Authors:** Adam J.E. Freedman, Kyle C. Peet, Jason T. Boock, Kevin Penn, Kristala L. J. Prather, Janelle R. Thompson

## Abstract

Supercritical carbon dioxide (scCO_2_) is an attractive substitute for conventional organic solvents due to its unique transport and thermodynamic properties, its renewability and labile nature, and its high solubility for compounds such as alcohols, ketones and aldehydes. However, biological systems that use scCO_2_ are mainly limited to *in vitro* processes due to its strong inhibition of cell viability and growth. To solve this problem, we used a bioprospecting approach to isolate a microbial strain with the natural ability to grow while exposed to scCO_2_. Enrichment culture and serial passaging of deep subsurface fluids from the McElmo Dome scCO_2_ reservoir in aqueous media under scCO_2_ headspace enabled the isolation of spore-forming strain *Bacillus megaterium* SR7. Sequencing and analysis of the complete 5.51 Mbp genome and physiological characterization revealed the capacity for facultative anaerobic metabolism, including fermentative growth on a diverse range of organic substrates. Supplementation of growth medium with L-alanine for chemical induction of spore germination significantly improved growth frequencies and biomass accumulation under scCO_2_ headspace. Detection of endogenous fermentative compounds in cultures grown under scCO_2_ represents the first observation of bioproduct generation and accumulation under this condition. Culturing development and metabolic characterization of *B. megaterium* SR7 represent initial advancements in the effort towards enabling exploitation of scCO_2_ as a sustainable solvent for *in vivo* bioprocessing.

## Introduction

Academic and industrial research has increasingly focused on supercritical carbon dioxide (scCO_2_) as an environmentally benign substitute for conventional organic solvents that are typically hazardous, toxic, and/or flammable (Budisa and Schulze-Makuch, 2014; Knez et al., 2014), including for use in broad classes of *in vitro* biocatalysis reactions that are challenging or expensive in aqueous phase reactors (Nakamura et al., 1986; Marty et al., 1992; Matsuda et al., 2004; Salgın et al., 2007; Matsuda et al., 2008; Wimmer and Zarevucka, 2010). Due to its unique thermodynamic and transport properties, scCO_2_ has been explored for a variety of bioprocessing applications that are expertly reviewed in the following: (Ikushima, 1997; Hobbs and Thomas, 2007; Darani and Mozafari, 2009; Matsuda, 2013). Additionally, scCO_2_ has been found to stabilize enzymes (Matsuda, 2013; Budisa and Schulze-Makuch, 2014), providing further benefits to *in vitro* reactions using this solvent. Downstream processing utilizing scCO_2_ has focused on extraction of compounds such as alcohols, lipids and acids from fermentation broth, as well as coffee decaffeination (Darani and Mozafari, 2009). Due to its capacity to disrupt the cellular membranes of a variety of microorganisms (Sun et al., 2006; Darani and Mozafari, 2009; Sahena et al., 2009), scCO_2_ is also commonly used as a sterilizing agent, especially in the food industry, where stainless steel piping and consumable products are difficult or undesirable to autoclave.

Microbial growth within or exposed to organic solvents has been discovered, described and engineered (Sardessai and Bhosle, 2002; Budisa and Schulze-Makuch, 2014; Mukhopadhyay, 2015); however, active growth in the presence of scCO_2_ has been rarely demonstrated. The rapid, sterilizing effects of scCO_2_ on nearly all bacteria due to cellular membrane disruption, desiccation, enzyme inactivation and cytosolic acidification (Spilimbergo and Bertucco, 2003) have rendered its use ineffective for *in vivo* biocatalysis in growing cultures (Knutson et al., 1999; Khosravi-Darani and Vasheghani-Farahani, 2005). Despite toxicity issues, microbial growth in the presence of scCO_2_ would offer multiple advantages over *in vitro* processes by enabling cellular protection and regeneration of catalytic enzymes and cofactors for *in vivo* pathways. In addition, enzyme secretion from growing cells would allow for the continuous generation of catalyst, and could enhance chemistries available under scCO_2_, such as cellulose deconstruction (Yang et al., 2015). Due to the solvent properties of scCO_2_, it could also be used *in situ* to extract toxic intermediates that may inhibit host growth or to harvest valuable bioproducts, reducing the need for downstream purification (Tompsett et al., 2018). Temperatures and pressures under which scCO_2_ forms (>31.1°C and >72.9 atm) are also compatible with the culture of barotolerant mesophilic organisms. Since scCO_2_ is lethal to most microorganisms, culture of biocompatible strains under scCO_2_ would likely resist contamination by off-target organisms, thus eliminating the need for costly antibiotics or specialized feeding strategies (Shaw et al., 2016). Lastly, culturing of cells under scCO_2_ is being investigated to enhance the efficacy of geological CO_2_ sequestration strategies through microbially-facilitated processes, including carbonate biomineralization and biofilm production to mitigate CO_2_ plume leakage (Mitchell et al., 2008; Peet et al., 2015).

Several studies have demonstrated the use of scCO_2_ as a reagent for biologically-catalyzed single-step reactions, including application to immobilized *Geotrichum candidum* cells (Matsuda et al., 2000) for ketone reduction and *Bacillus megaterium* cells for pyrrole carboxylation (Matsuda et al., 2001). However, in both of these cases, the cell material was not grown under scCO_2_, the reaction time period was constrained to several hours, and the cellular viability and activity of multi-step metabolic pathways under scCO_2_ was not established. To be considered feasible for industrial bioprocess development, it is required for strains to remain biocompatible with scCO_2_ in a batch or continuous flow bioreactor over a time period of days to weeks. In light of this requirement, recent attention has focused on several research efforts that have identified microbial populations resilient over extended periods of exposure to scCO_2_ in natural (Mu et al., 2014; Freedman et al., 2017) and laboratory (Mitchell et al., 2008) systems, including in aqueous media under scCO_2_ headspace (Peet et al., 2015).

Inspired by the identification of scCO_2_-resistant bacterial communities, especially those detected in natural environments, we sought to discover and develop a scCO_2_ biocompatible laboratory strain that would enable routine culture and bioproduction. Previous work attempting to establish strains able to reliably grow under scCO_2_ proved challenging. Individual cultures started from *Bacillus* spores in steel columns under scCO_2_ had growth frequencies below 50% over a period of several weeks (Peet et al., 2015). Building on the scCO_2_ enrichment and culturing protocols developed in this previous work, the current study sought to improve growth outcomes under scCO_2_ by utilizing a bioprospecting approach that targeted environmental microorganisms exposed to scCO_2_ over geologic time scales. We thus collected fluid samples sourced from the deep subsurface at McElmo Dome CO_2_ Field in Cortez, Colorado, where scCO_2_ has accumulated over 40-72 million years (Cappa and Rice, 1995; Gilfillan et al., 2008). Since previous work has revealed evidence of an anaerobic microbial ecosystem (Freedman et al., 2017) we hypothesized that organisms isolated from this site may demonstrate enhanced scCO_2_-resistance and growth potential.

We report isolation of a *Bacillus megaterium* strain, denoted SR7, capable of consistent and robust growth by fermentative metabolism in pure culture under our laboratory scCO_2_ conditions. *B. megaterium* SR7 was genomically and physiologically characterized to establish optimal culturing conditions for growth with metabolite production under scCO_2_. We also report the use of chemical induction of endospore germination to enable significant improvement in SR7 growth frequency under scCO_2_. Successful demonstration of this unique culturing concept is a significant advancement in the development of *in vivo* bioproduction using metabolically active wild-type cells exposed to the sustainable solvent scCO_2_.

## Materials and Methods

### Supercritical CO_2_-exposed enrichment culture

#### Enrichment cultures and serial passaging

Formation water from the McElmo Dome CO_2_ Field sourced from deep subsurface CO_2_ production wells within the Yellow Jacket and Hovenweep Fields (operated by KinderMorgan CO_2_) was collected from fluid-gas separators that were decanted and filled 15 hours prior to sampling. One liter of degassed fluid from each well was collected in an acid-washed bottle (Nalgene), shipped on ice and stored at 4°C until used as enrichment culture inocula. 100 ml of fluids from three wells (HB-5/Well 2, HE-1/Well 4, YB-4/Well 7) that contained biomass as observed by epifluorescence microscopy (Freedman et al., 2017) were filtered onto 0.22 μm polycarbonate filters (Whatman Nucleopore) to concentrate microbial cells. Using sterilized tweezers within an anaerobic chamber, filters were placed inside high-pressure culturing vessels with 1 ml of formation water from the same well, and 4 ml of growth medium. The McElmo Dome enrichments were incubated under scCO_2_ for 45 days (first incubation, designated M1), after which enrichments inspected by epifluorescence microscopy showing biomass accumulation were serially passaged by dilution in degassed growth medium to starting concentrations of 10^4^ cells/ml. Subsequent rounds of passaged cultures were incubated under scCO_2_ for 19 days (M2), 33 days (M3), and 35 days (M4).

#### Culturing media and vessels

Due to elevated sulfur content and detection of DNA from heterotrophic and sulfate reducing bacteria in McElmo Dome fluids (Freedman et al., 2017), media for enrichment culture and serial passaging was a modified version of sulfate-amended rich MS medium (Colwell et al., 1997), denoted MS-SR medium, as described in Peet et al. (2015). MS-SR medium consisted of (in g/l) 0.5 yeast extract, 0.5 tryptic peptone, 10.0 NaCl, 1.0 NH_4_Cl, 1.0 MgCl_2_•6H_2_O, 0.87 K_2_SO_4_, 0.83 FeSO_4_, 0.82 sodium acetate, 0.4 K_2_HPO_4_, 0.4 CaCl_2_, 0.0025 EDTA, 0.00025 CoCl_2_•6H_2_O, 0.0005 MnCl_2_•4H2O, 0.0005 FeSO_4_•7H_2_O, 0.0005 ZnCl_2_, 0.0002 AlCl_3_•6H_2_O, 0.00015 Na_2_WoO_4_•2H_2_O, 0.0001 NiSO_4_•6H_2_O, 0.00005 H_2_SeO_3_, 0.00005 H_3_BO_3_, and 0.00005 NaMoO_4_•2H_2_O. Luria-Bertani Broth (LB; Difco) was included as an additional growth medium for the final passage (M4). All media used in scCO_2_ passaging was amended with 0.25 g/l of reducing agent Na_2_S and 0.001 g/l of the redox indicator resazurin. A summary of media formulations is presented in Table S1.

High-pressure vessels for culturing under scCO_2_ were constructed of ¾ inch 316 stainless steel tubing for 10 ml total capacity, and fitted with quarter turn plug valves (Swagelok or Hylok). Between uses, all vessel and pressurization manifold components were cleaned and sterilized as described in Peet et al., (2015). Prior to vessel loading, culture medium was degassed with a stream of extraction grade CO_2_ for 30 minutes. Vessels were then filled to ½ capacity (5 ml) with inocula and degassed medium, after which the headspace was pressurized with CO_2_ gas at a rate of 2-3 atm/min until reaching a final pressure of 100 atm. After pressurization, reactors were incubated at 37°C shaking at 100 rpm. Prior to depressurization, vessels were connected via 316 stainless steel tubing and fittings to a gauge (Hunter) to verify that vessel pressures remained above the critical point. After depressurizing at a rate of 3-5 atm/min, vessels were transferred to an anaerobic chamber (Coy Lab Products; 95% CO_2_, 5% H_2_) for sampling, glycerol stock preparation and passaging into fresh medium.

#### Enumeration of cell density

To quantify cultured biomass, 0.5-1.0 ml samples were treated with Syto9 nucleic acid stain (Life Technologies), incubated in the dark for 15 minutes, collected on 0.22 μm polycarbonate filters (Whatman Nucleopore) by vacuum pump, washed twice with phosphate buffered saline (PBS), and mounted on glass slides with immersion oil and a cover slip for visualization by epifluorescence microscopy (Zeis Axioscope). Filters containing stained cells were stored at 4°C in the dark until use. Cell densities were extrapolated using individual cell counts as described in Peet et al. (2015). Final reported cell densities represent averaged cell counts in 15-20 separately viewed fields per sample. The limit of detection was considered to be half of one cell per 15 fields, which corresponds to 1.15×10^3^ cells/ml. Fluorescence images were captured on a Nikon D100 camera using the NKRemote live-imaging software. To verify media sterility, cell-free negative control incubations were inspected for biomass accumulation by fluorescence microscopy and overnight LB agar plating at 37°C.

### Strain isolation

Samples from the final round of passaging (M4) were plated on LB agar and incubated at 37°C under aerobic conditions. Colonies were used to inoculate overnight liquid LB cultures to prepare glycerol stocks stored at −80°C. Genomic DNA extracted from overnight cultures using the Qiagen Blood and Tissue Kit (Gram-positive protocol) was used as template for 16S rRNA PCR using universal bacterial primers 515F (GTGCCAGCMGCCGCGGTAA) and 1492R (GGTTACCTTGTTACGACTT) (Turner *et al.*, 1999). PCR mixtures included 1X Phusion High Fidelity Polymerase buffer, 0.4 μM of each primer (IDT), 0.4 μM deoxynucleotide mixture and 1 U Phusion Polymerase (New England Biolabs). PCR products were purified using Exo-SAP IT (Affymetrix) and submitted for Sanger sequencing (Genewiz). Primer removal, universal trimming, sequence alignment and tree building using a bootstrapped (100X) neighbor-joining method was conducted in CLC Genomics Workbench Version 7 (Qiagen), with tree visualization in FigTree v1.4.2. 16S rRNA gene reference sequences were downloaded from Genbank (NCBI).

### Spore preparation

*B. megaterium* SR7 was prepared as spores using the protocol described in Peet et al. (2015), based on the original method developed in Kim and Goepfert (1974). Spore preparations were heat-treated at 80°C for 10 min to inactivate residual vegetative cells, stored in wash buffer at 4°C until use, and tested for viability by counting colony forming units after plating on LB agar.

### Genome sequencing and analysis of isolate SR7

Isolate strain *B. megaterium* SR7 genomic DNA was extracted from a 10 ml overnight aerobic LB culture using the Qiagen Blood and Tissue Kit (Gram-positive protocol). Eluted DNA was submitted to the MIT BioMicro Center for sequencing using PacBio SMRT technology. PacBio assembler software was used to assemble SR7 contigs, which were compared to the genome of closely related strain *B. megaterium* QM B1551 (Eppinger et al., 2011) using the online tools NUCmer (Delcher et al., 2002) and Double Act (Carver et al., 2005), enabling an inter-genome Blastn comparison viewable in the Artemis Comparison Tool (ACT). Based on the ACT comparison, the SR7 chromosome was adjusted to start at the beginning of gene *dnaA* in agreement with the reference genome. The closed chromosome was then visualized by DNA Plotter (Carver et al., 2009) and submitted to Rapid Annotation using Subsystem Technology (RAST) (Overbeek et al., 2014) for open reading frame (ORF) gene prediction and functional annotation. Contig sequences that did not overlap with the assembled genome were classified as plasmids and submitted to RAST and Blastp for functional annotation. Shared and unique gene annotations between SR7 and *B. megaterium* reference genomes QM B1551, DSM319 (Eppinger et al., 2011), and WSH-002 (Liu et al., 2011) were elucidated using online tool Venny 2.1 (Oliveros, 2007-2015). Inter-strain sequence comparisons were conducted using the Average Nucleotide Identity (ANI) calculator (Rodriguez and Konstantinidis, 2016).

The SR7 genome and putative plasmids were searched against the virulence factor database core data set A (Chen et al., 2016) using Blastn. Any elements showing a high degree of sequence similarity (E-values <0.01) were flagged as possible toxins or virulence factors in SR7; some elements displaying E-values >0.01 were subjected to further investigation when potentially part of a system with multiple components showing high similarity to SR7 genes. Any identified potential risk factors that were not known to be common *Bacillus* toxins or virulence markers but verified to be part of annotated open reading frames in SR7 were compared to similar genes found in other strains of *B. megaterium* and *B. subtilis ssp. subtilis 168*.

### Physiological characterization of isolate SR7

Growth dynamics of isolate SR7 were first investigated under aerobic ambient pCO_2_ and anaerobic 1 atm CO_2_ anaerobic conditions due to their higher throughput and experimental simplicity relative to scCO_2_ incubations. 1 atm CO_2_ is also considered a proxy for pressurized conditions due to its intermediate pH and predicted dissolved CO_2_ concentration (pH 5, 2.6×10^−2^ M) relative to ambient air (pH 7, 1.2×10^−5^ M) and scCO_2_ (pH 3.5, 2.7 M) (Peet et al., 2015). To determine growth of SR7 with respect to pH, salinity, and bicarbonate concentration under aerobic ambient pCO_2_ conditions, 10^4^ spores/ml were inoculated in 5 ml LB medium and incubated on a spinning rack at 100 rpm for 24 hours at 37°C. pH levels (pH 2-10) were titrated with HCl or NaOH while the effect of salinity or bicarbonate on growth was determined by adding NaCl (1-10%) or NaHCO_3_ (0.1-0.5 M), respectively. SR7 growth was assayed over a temperature range of 9-55°C. Antibiotic sensitivity was determined by supplementing 5 ml LB medium with ampicillin (5-50 μg/ml), chloramphenicol (3.5-35 μg/ml), kanamycin (5-50 μg/ml), spectinomycin (10-100 μg/ml), streptomycin (10-100 μg/ml), or tetracycline (1.5-15 μg/ml) (additional sensitivities were assessed using Biolog GenIII plate cultures, as described below). Conditional growth effects and antibiotic sensitivities were determined by comparing OD_600_ measurements of amended cultures to unamended positive and cell-free negative LB medium controls. Efficacy of sterilization methods were assayed by plating spores or cell preparations on LB agar medium before and after treatment with 10% bleach for 20 minutes or autoclaving at 121°C for 35 minutes to determine the change in viability associated with treatment.

Minimal medium for SR7 consisted of a M9 salt base amended with 4 g/l glucose or 4 g/l xylose as sole carbon sources, with or without trace metals solution (Boone et al., 1989). The 1X concentration trace metals solution consisted of (in mg/l): 5 Na_2_(EDTA), 0.2 NiSO_4_•6H_2_O, 0.5 CoCl_2_•6H_2_O, 0.1 H_2_FeO_3_, 1 FeSO_4_•7H_2_O, 0.1 H_3_BO_3_, 1 ZnCl_2_, 0.1 NaMoO_4_•2H_2_O, 0.4 AlCl_3_•6H_2_O, 1 MnCl_2_•4H_2_O, 0.3 Na_2_NO_4_•2H_2_O, 0.2 CaCl_2_. Dilute LB (0.001-0.01X) medium or yeast extract (YE; 5-50 mg/l) were used as vitamin/co-factor supplements for M9+ medium (Table S1), and 5 mM NaNO_3_ was included as a potential alternative electron acceptor for a subset of media combinations where specified.

SR7 cultures incubated under 1 atm CO_2_ anaerobic headspace were grown in 10 ml of CO_2_-degassed medium (LB or M9+) in 100 ml serum vials with clamped rubber stoppers at 37°C. Anaerobic cultures were inoculated with spores (10^4^ spores/ml) or vegetative cells passaged under anaerobic conditions. For passaged vegetative cells, an overnight anaerobic culture of SR7 was sub-cultured in at least triplicate to OD_600_ of 0.01 in minimal (M9+; Table S1) or LB medium. Growth under 1 atm CO_2_ was assessed over 28-32 hours by measurement of OD_600_ via 96-well microplate reader (BioTek Synergy 2) or 1 cm pathlength nanophotometer (Implen), colony forming units (CFUs) on LB agar plates, and glucose consumption (HPLC, Agilent 1100 Series). Doubling times were calculated using a log-linear fit of OD_600_ data points during exponential growth. Shake speeds of 150, 250, and 350 rpm were tested to determine the optimal disruption regime for anaerobic 1 atm CO_2_ cultures based on doubling rate and prolonged culture viability.

Biolog GenIII (Biolog; Hayward, CA) (Bochner, 2003) 96-well plates were used to assess single carbon source utilization and antibiotic/chemical sensitivities for SR7 and other *B. megaterium* strains under aerobic conditions. Colonies grown overnight on solid BUG medium (Biolog) were added to Inoculating Fluid B (Biolog) to a percent transmittance of 90-94% at 600 nm. For some samples, the inoculating fluid was amended with 1X trace metals solution (Boone et al., 1989). Upon bacterial colony addition, inoculating fluid was transferred to a GenIII plate, which was incubated at 37°C on a plate shaker at 800 rpm. Growth in each well was quantified using colorimetric changes measured by OD_490_ on a microplate reader (BioTek Synergy 2) and integrating the area under the curve of OD_490_ values over the course of the incubation at standardized time points. Growth on each substrate and antibiotic was categorized as follows: “−” displays an area less than the negative control, “+” is greater than the negative control, but less than half of the positive control, while “+++” is between “+” and the positive control (grown in rich medium).

### Imaging by scanning electron microscopy (SEM)

Spore preparations and vegetative cell cultures grown under aerobic, 1 atm CO_2_, and scCO_2_ conditions were collected on 0.22 μm polycarbonate filters (Whatman Nucleopore) by vacuum pump. Filters were then immersed for 4 h in 40 g/l paraformaldehyde. Fixed cells were washed once with PBS, dehydrated with increasing concentrations of ethanol (50, 75, 90, 95, 100, 100, 100%) over the course of 2 hours, then stored overnight at 4°C. Residual ethanol was subsequently removed by immersion in supercritical carbon dioxide. After critical point drying, filters were sliced into smaller sections, placed on stubs, and coated with gold/palladium using the Hummer 6.2 Sputter System. Images were captured using a Jeol 5600LV scanning electron microscope operating at a working distance between 10-25 mm and acceleration voltage of 5 kV.

### Analysis of SR7 fermentation products under 1 atm CO_2_ and scCO_2_

SR7 cultures incubated under 1 atm CO_2_ or scCO_2_ in M9+ or LB medium were centrifuged for 5 minutes at 21,000 x g, and the cell-free supernatant was collected. Glucose consumption and extracellular fermentation products were analyzed by HPLC (Agilent 1100 Series). Supernatants were loaded onto an Aminex HPX-87H anion exchange column (Bio-Rad, Hercules, CA) using 5 mM H_2_SO_4_ as the mobile phase with a flow rate of 0.6 ml/min, and a column inlet and outlet temperature of 35°C. Refractive index, area under the curve, and standard curves for analytes (including retention time) were used to identify and quantify glucose consumption and fermentation product generation.

### Evaluating prospective spore germination inducers

Two amino acids (L-alanine and L-leucine) known to facilitate germination through independent mechanisms (Hyatt and Levinson, 1962), as well as heat activation treatment, were selected for investigation with SR7. Triplicate cultures in 10 ml degassed LB medium were inoculated with SR7 spores to an OD_600_ of 0.01 in serum vials. LB medium was amended with 100 mM L-alanine or subjected to heat activation by placement in a 65°C water bath for 15 minutes after inoculation; unamended LB medium was used as a control. Cultures grown at 37°C while shaking at 250 rpm were analyzed for growth at multiple time points by measuring OD_600_ and colony counts on LB agar medium. After plating each time point, samples were exposed to 80°C for 10 minutes to heat kill vegetative cells (Setlow, 2006), followed by plating on solid LB medium to ascertain the remaining number of spores.

To decouple the spore germination process from vegetative cell outgrowth, germination inducers were tested in buffered media without nutrient sources. SR7 spores were added in triplicate to 10 ml degassed PBS in serum vials amended with 100-250 mM L-alanine, 25 mM L-leucine, or unamended PBS as a control. A second set of 100 mM L-alanine cultures was heat activated by incubating at 65°C for 15 minutes post-inoculation. Amino acid amendment concentrations and heat activation parameters were chosen based on maximum germination induction rates shown in previous studies (Hyatt and Levinson, 1962; Levinson and Hyatt, 1970). PBS suspensions were incubated at 37°C while shaking at 250 rpm for 9 hours. Time point samples (1 ml) were stained with Syto9, filter collected and prepared for epifluorescence microscopy, as previously described.

The proportion of germinated spores was determined by epifluorescence microscopy using the staining method validated by Cronin and Wilkinson (2007). 100-300 spores were counted per filter and categorized as either “dormant” or “germinated” depending on whether the stain was localized to the cell membrane or diffused within the interior of the cell, respectively. Syto9 localizes to the exterior of the cell in dormant spores due to its inability to penetrate the spore’s protective outer coat, while germinated cells appear completely stained as Syto9 diffuses into the cell upon spore coat degradation (Cronin and Wilkinson, 2007). Cells displaying an intermediate degree of stain dispersal were categorized as germinated. Due to the increased extent of emitted fluorescence in germinated cells, bulk fluorescence was an additional metric used to quantify population-wide germination. Bulk fluorescence (485/20 excitation, 528/20 emission) was measured on a microplate reader (BioTek Synergy 2).

### Evaluating SR7 scCO_2_ growth under optimized conditions

SR7 growth outcomes under scCO_2_ headspace (90-100 atm, 37°C) were investigated using conditions and media formulations informed by characterization under aerobic and 1 atm CO_2_ (including shaking at 250 rpm). LB medium was amended with 50 mM disodium phosphate (P-LB) to standardize buffering capacity and phosphate content with M9+ medium. To examine the influence of germination inducers, SR7 spores inoculated at starting concentrations of 3×10^4^ spores/ml in 5 ml of P-LB or M9+ medium were amended with either 100 mM L-alanine (M9A+), 10 mM L-leucine (P-LBL; Table S1), or were heat activated at 65°C for 15 minutes prior to 37°C incubation (M9+ cultures only). After incubation under scCO_2_ for 18-20 days, growth was identified in depressurized cultures by observation of vegetative cell morphologies using fluorescence microscopy, and an increase of >10-fold cell counts relative to loaded spore concentrations. To determine the statistical significance of germination inducers or media type, a non-parametric Wilcoxon/Kruskal-Wallis Test was performed on the dataset (JMP Pro v.12) where outcome (growth/no growth) or cell density fold change (relative to t_0_) were dependent variables and incubation time and L-alanine presence/absence were independent variables.

To establish whether increasing starting spore concentrations and incubation times improve the likelihood of growth, replicate SR7 cultures in M9A+ medium loaded with four starting spore concentrations (5.6×10^5^, 4.6×10^3^, 5×10^1^, 5×10^−1^ cells/ml) were incubated over an 18-day time course and scored for growth by filter cell counts. Because reactors inoculated with 5×10^1^ and 5×10^−1^ cells/ml are below the limit of detection, their concentrations were recorded as one half the detection limit (1.15×10^3^ cells/ml). Time course data was combined with prior scCO_2_ incubation results in identical M9A+ medium to develop a logistic regression model (JMP Pro v. 12) for growth frequency where outcome (growth/no growth) was the dependent variable, and inocula concentration and incubation time were independent variables.

## Results

### Isolation of scCO_2_-biocompatible strain SR7 from McElmo Dome formation fluid

Fluids from three CO_2_ production wells at McElmo Dome, previously shown by microscopy to contain biomass (Freedman et al., 2017), were used as inocula for enrichment culture under scCO_2_. The well-fluid enrichment cultures incubated under 100 atm CO_2_ pressure for 45 days (M1) revealed a majority of cultures with cell concentrations greater than 10^6^ cells/ml (Table 1A), likely reflecting filter-captured biomass used to inoculate M1 reactors (see methods; the predicted inoculum concentration for Well 7 fluids (3.8×10^6^ cells/ml) is approximately equal to the post-M1 incubation cell count (3.0×10^6^ cells/ml)). To isolate individual strains capable of growth, reactors containing the highest final cell concentrations were diluted >100-fold to 10^4^ cells/ml and serially passaged into fresh culture medium for three subsequent rounds of incubation (M2-M4; Table 1A). Passaged cultures originally inoculated with fluids from Well 7 demonstrated repeated biomass accumulation over all four rounds of passaging, with maximum cell densities in the final passage (M4) observed to be 6.9×10^6^ and 7.8×10^6^ cells/ml in MS-SR and LB media, respectively.

**Table 1.**
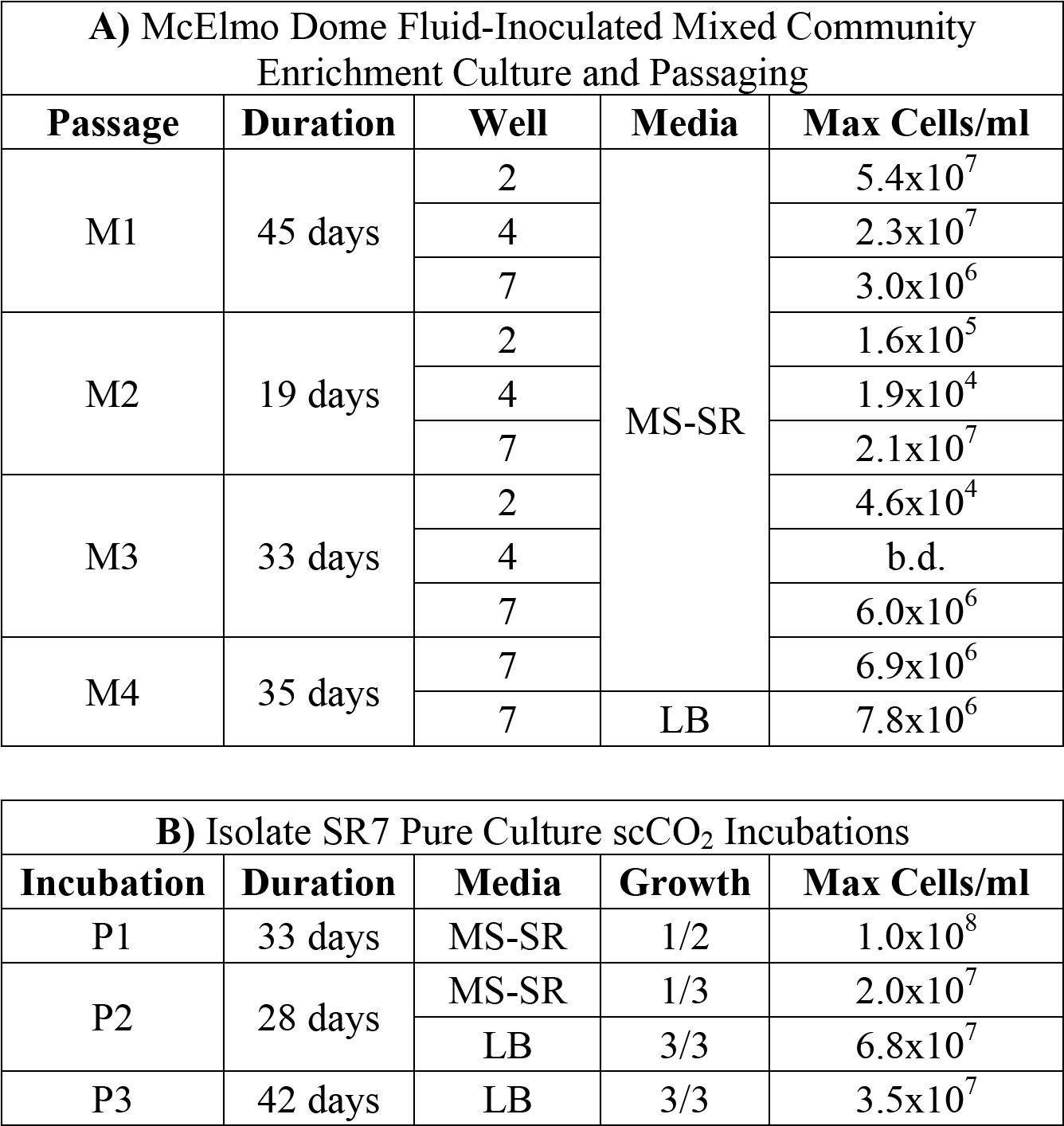
Summary of scCO_2_-resistant strain isolation results

| A) McElmo Dome Fluid-Inoculated Mixed Community Enrichment Culture and Passaging |  |  |  |  |
| --- | --- | --- | --- | --- |
| Passage | Duration | Well | Media | Max Cells/ml |
| M1 | 45 days | 2 | MS-SR | 5.4x10<sup>7</sup> |
|  |  | 4 |  | 2.3x10<sup>7</sup> |
|  |  | 7 |  | 3.0x10<sup>6</sup> |
| M2 | 19 days | 2 |  | 1.6x10<sup>5</sup> |
|  |  | 4 |  | 1.9x10<sup>4</sup> |
|  |  | 7 |  | 2.1x10<sup>7</sup> |
| M3 | 33 days | 2 |  | 4.6x10<sup>4</sup> |
|  |  | 4 |  | b.d. |
|  |  | 7 |  | 6.0x10<sup>6</sup> |
| M4 | 35 days | 7 |  | 6.9x10<sup>6</sup> |
|  |  | 7 | LB | 7.8x10<sup>6</sup> |

| B) Isolate SR7 Pure Culture scCO<sub>2</sub> Incubations |  |  |  |  |
| --- | --- | --- | --- | --- |
| Incubation | Duration | Media | Growth | Max Cells/ml |
| P1 | 33 days | MS-SR | 1/2 | 1.0x10<sup>8</sup> |
| P2 | 28 days | MS-SR | 1/3 | 2.0x10<sup>7</sup> |
|  |  | LB | 3/3 | 6.8x10<sup>7</sup> |
| P3 | 42 days | LB | 3/3 | 3.5x10<sup>7</sup> |

Since cultures inoculated with fluids from Wells 4 and 2 were unable to maintain biomass viability after passages M2 and M3, respectively, strain isolation efforts focused exclusively on passages originally inoculated with fluids from Well 7. The isolation of a single dominant strain from Well 7 passages was confirmed by uniform cell (Figures 1A-B) and colony morphology, as well as 16S rRNA gene sequencing, which classified the isolate as *Bacillus megaterium* or *Bacillus aryabhattai* based on sequence homology. A 16S rRNA gene-based phylogenetic tree (Figure 2) shows a high degree of similarity between SR7 and other well-established subsurface and scCO_2_-associated species (Freedman et al., 2017; Peet et al., 2015) and genomic analysis allowed tentative assignment of the isolate to the species *B. megaterium* (see Genomic Analysis section). Microscopic inspection of M4-passaged cultures grown under scCO_2_ revealed both rod shaped vegetative cells and spherical endospores (Figures 1A-B; vegetative cells shown).

**Figure 1.**
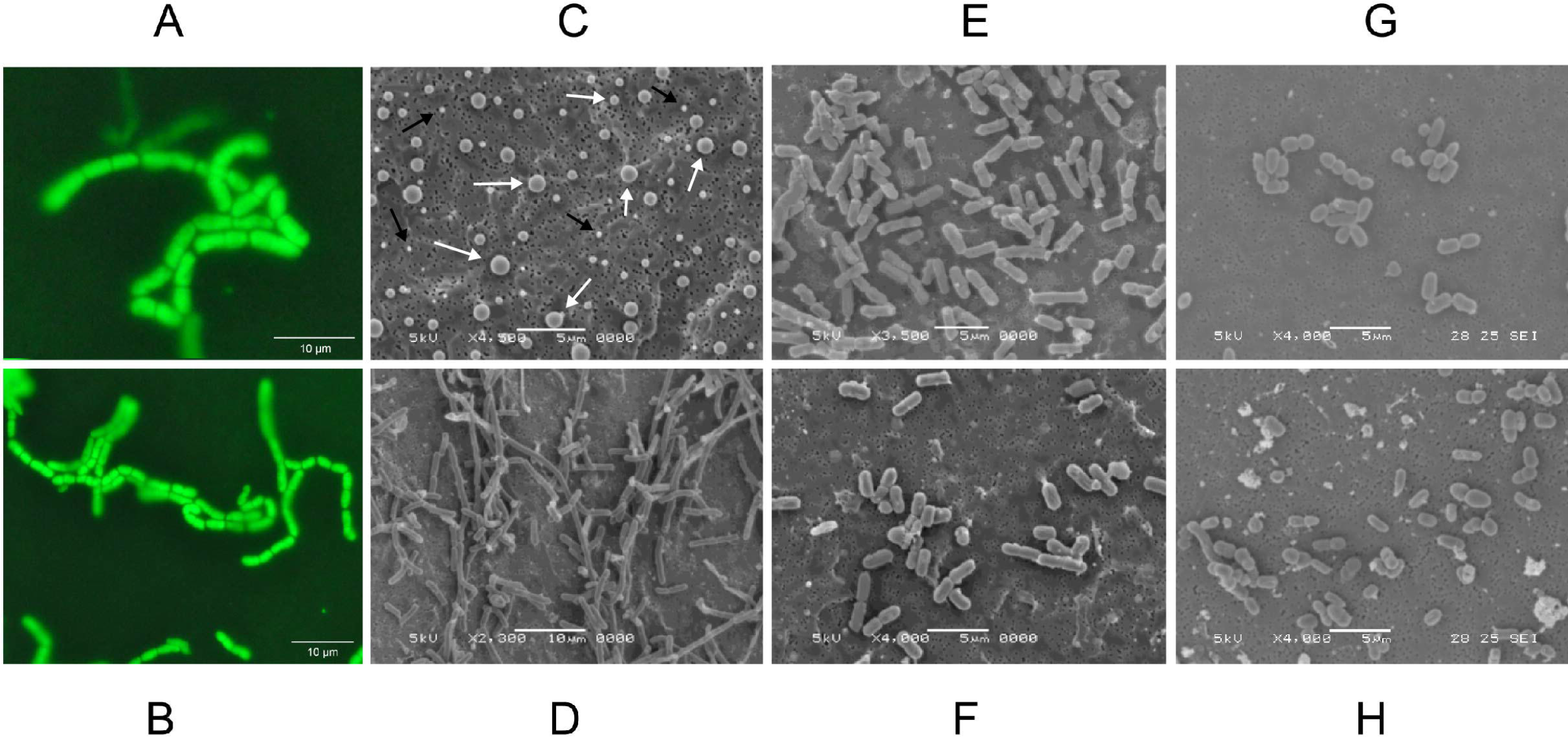
**A-B)** Fluorescence microscopy images of *B. megaterium* SR7 vegetative cells grown during scCO_2_ passage M4 in MS-SR medium (multiple images of the same incubation at different magnification). Cells were treated with Syto9 nucleic acid stain for fluorescence visualization. Scanning electron microscope (SEM) images of *B. megaterium* SR7 **C)** prepared endospores of varying size, as indicated by white arrows (and salt crystals indicated by black arrows), **D)** aerobic cultures in LB medium, **E)** aerobic cultures in M9+ medium, **F)** 1 atm CO_2_ cultures in M9+ medium, and **G-H)** scCO_2_ cultures grown in M9A+ medium.

Isolate *B. megaterium* SR7 was prepared as spores (Figure 1C) for long-term storage, and to confirm its ability to germinate and grow under scCO_2_, as previously demonstrated for other *Bacillus* strains (Peet et al., 2015). Spore preparations examined by SEM revealed a heterogenous distribution of sizes consistent with prior observations of various Bacillus species (Carrera et al., 2006; Leuschner et al., 1999; Hitchins et al., 1972) (Figure 1C). Final cell concentrations and elevated growth frequencies from mixed-culture scCO_2_ enrichments (M1-M4, Table 1A, Figures 1A-B) and spore-inoculated pure culture incubations (P1-P3; Table 1B), established *B. megaterium* SR7 as capable of consistent, robust growth under scCO_2_, especially in LB medium (e.g. 6/6 growth after 4 to 6 weeks of incubation in P2-P3; Table 1B). Spore preparations of *B. megaterium* SR7 maintained viability over 2 years at 4°C, suggesting that 4°C storage of SR7 spore stocks is appropriate if the strain is to be utilized for laboratory culturing.

**Figure 2.**
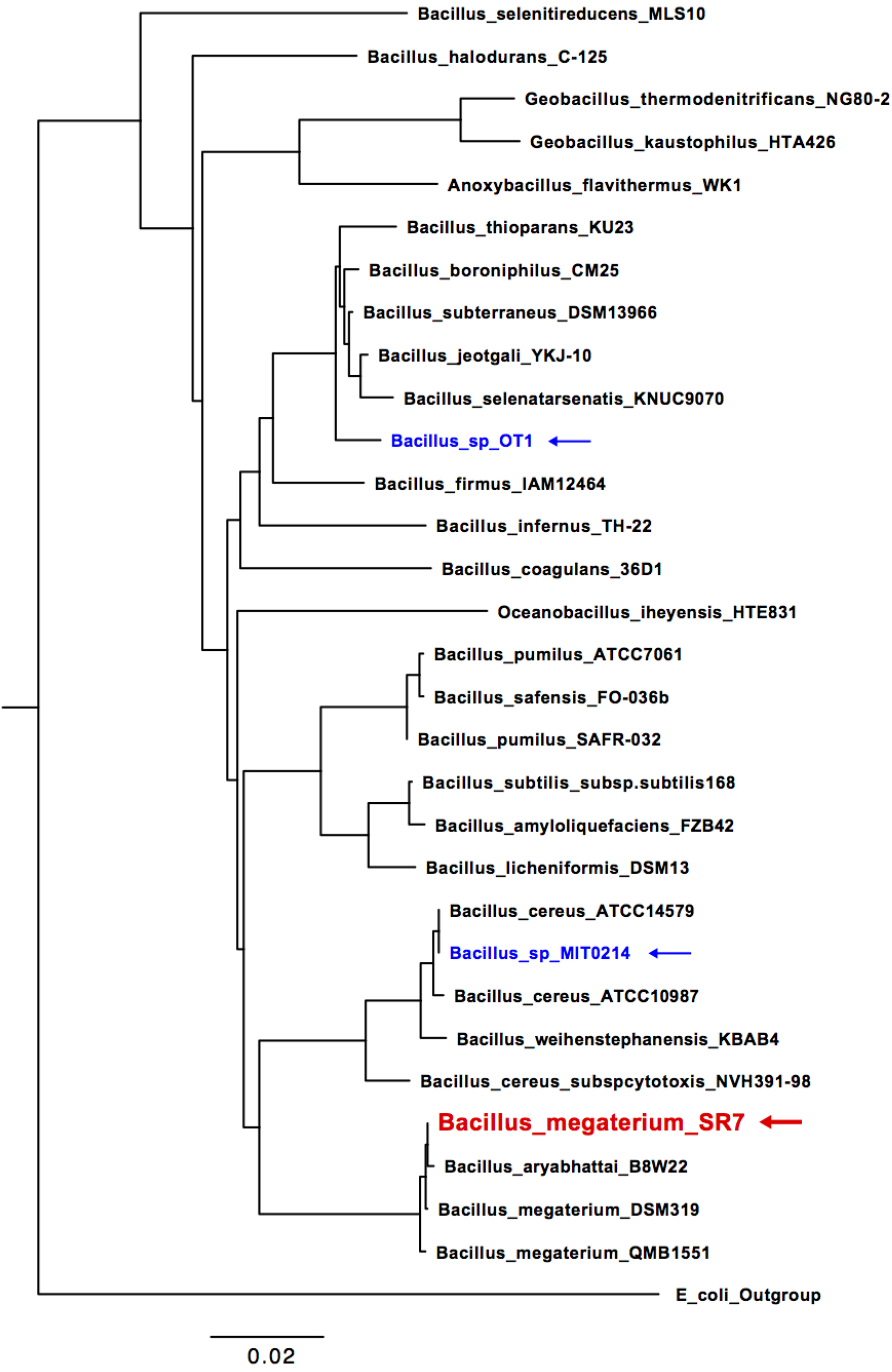
16S rRNA phylogenetic tree of McElmo Dome isolate *B. megaterium* SR7 (red), isolates investigated in Peet *et al*. (2015; blue) and additional closely related Bacilli (collected from NCBI GenBank). *E. coli* strain O91 was used as an outgroup.

### Physiological Characterization of SR7

We initiated strain characterization and culturing optimization to reduce SR7 scCO_2_ incubation times to <3 weeks in a semi-defined minimal medium while maintaining a high frequency of growth outcomes, increasing biomass accumulation, and generating fermentative products to verify and expand the metabolic capacity of SR7 under laboratory scCO_2_ conditions.

#### Safety assessment

Although *Bacillus megaterium* strains are considered biosafety level 1 organisms, we performed a safety assessment of new environmental isolate strain *B. megaterium* SR7 to ensure safe culturing and laboratory containment. Sterilization of vegetative *B. megaterium* SR7 cells was accomplished by standard 10% bleach treatment for 20 minutes, while autoclave treatment at 121°C was found to inactivate SR7 spores, with both techniques achieving >6-log reduction in viability (data not shown). After establishing safe disposal techniques, we aimed to assess the sensitivity of SR7 to an array of antibiotics on solid LB medium or using a panel of known toxic compounds at standard concentrations in a Biolog Gen III microplate (Table 2). We found SR7 to be sensitive to all antibiotics tested except spectinomycin, demonstrating safe measures for controlling SR7 growth, as well as establishing prospective antibiotic selection markers for genetic engineering. Compound sensitivity assays also revealed SR7 growth to be inhibited by D-serine and Niaproof 4, which are known to disrupt cell wall synthesis and lipid membranes, respectively.

**Table 2.**
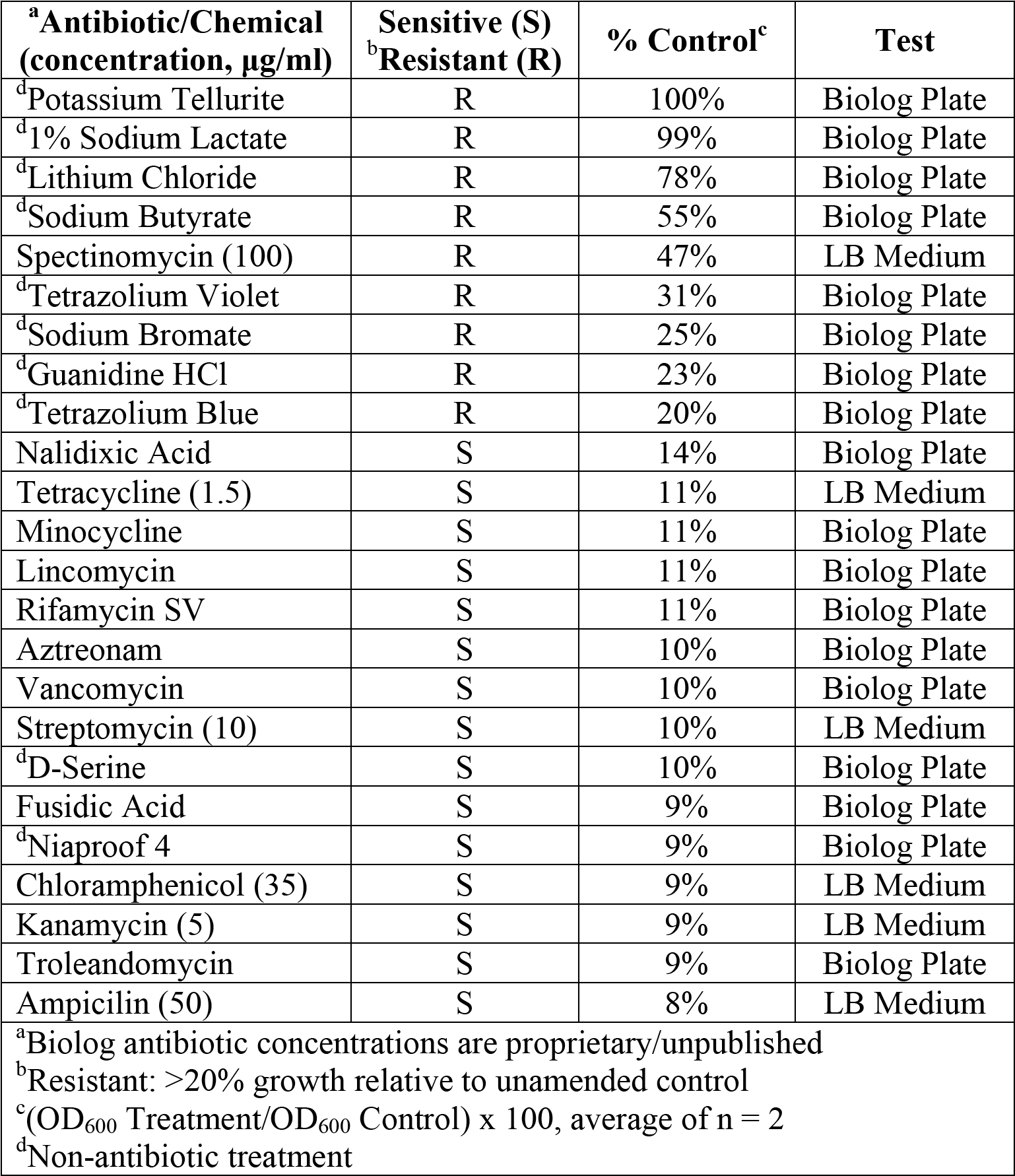
SR7 antibiotic/chemical sensitivity assay summary

#### Analysis of physiochemical variables affecting growth

Parameters influencing SR7 growth were first tested in aerobic and 1 atm CO_2_ anaerobic cultures because these experiments are shorter (1-3 days) than those under scCO_2_ (<3 weeks) and higher throughput due to ease of culturing setup (Table 3). Under aerobic conditions the optimal pH for SR7 growth was between 6 and 7, with growth observed from pH 4 to 10. Notably, the approximate media pH under scCO_2_ is between pH 4 and 5 depending on buffering capacity (Peet et al., 2015). SR7 tolerated bicarbonate concentrations up to 300 mM, with optimal growth below 100 mM; decreasing growth rate with increasing bicarbonate indicates an adverse response to concentrated dissolved CO_2_ species, and an upper limit on the positive effect of carbonate media buffering. Lastly, incubation temperature was assessed and growth found to be supported between 23-45°C, with optimal growth at 37°C, as commonly observed for other *Bacillus* species (Warth, 1978). Increased shake speeds over the tested interval (150-350 rpm) were observed to facilitate more rapid vegetative outgrowth from spores (Figures 3A-B) and passaged vegetative culture growth (Figures 3C-D) under 1 atm CO_2_ in LB medium. Cultures incubated at 350 rpm maintained high optical density while viable cell counts declined sharply after 18 hours (Figure 3C). An incubation shake speed of 250 rpm was utilized for subsequent incubation experiments to maximize doubling rates while sustaining viability into stationary phase.

**Figure 3.**
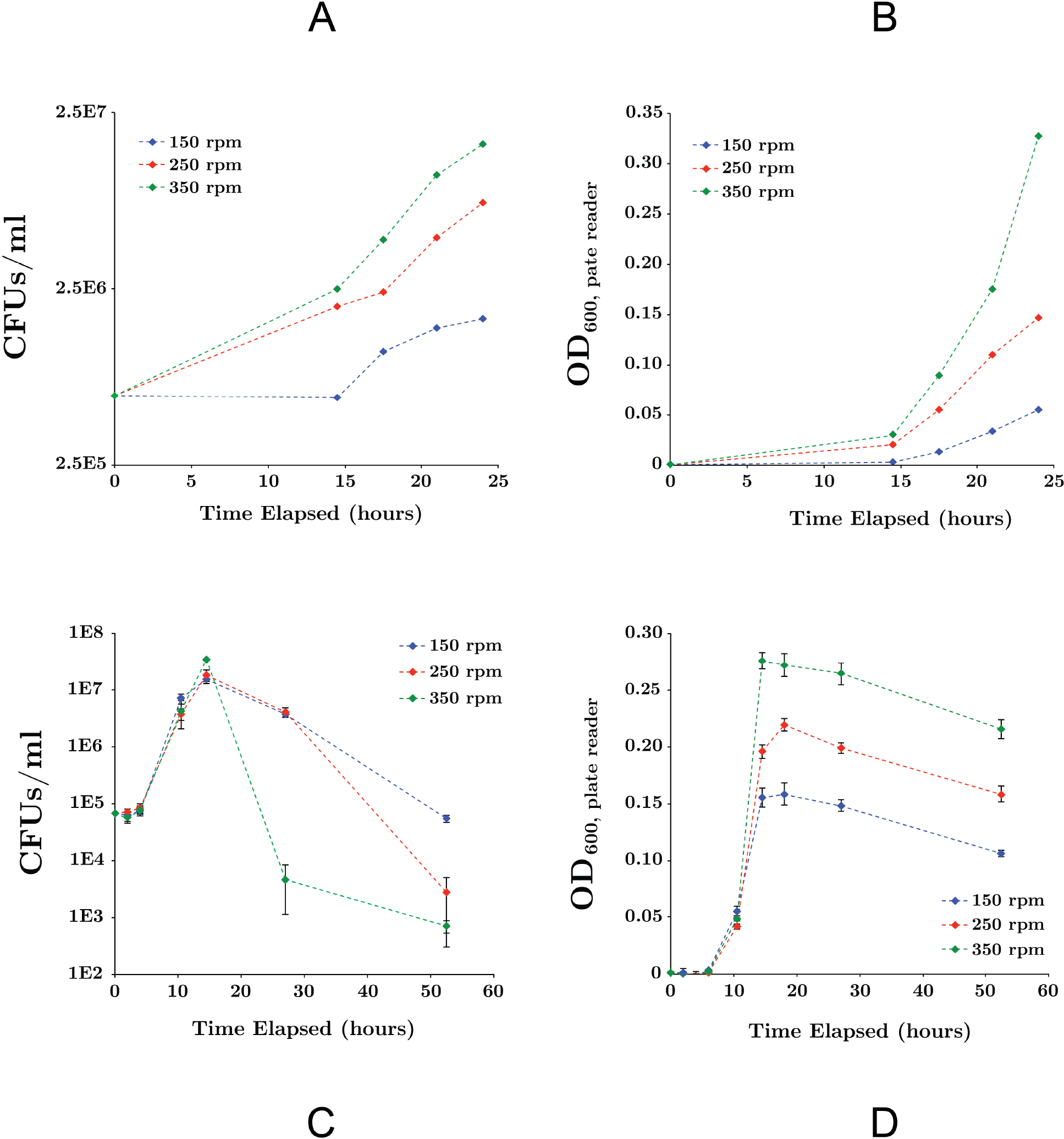
Effect of mixing rates on SR7 growth in LB medium under 1 atm CO_2_ for cultures inoculated from spores (A, B) or vegetative cells (C, D). Growth was measured by CFUs/ml **A)** and **C)** and OD_600_ **B)** and **D)**. Optical density was measured in 200 μl using a microplate reader and error bars represent the standard deviation of triplicate cultures. As expected, lag times for cultures inoculated with spores were longer than for those inoculated with vegetative cells due to time required for germination.

**Table 3.** Summary of viable SR7 aerobic growth in LB medium

| Condition | Range | Optimal |
| --- | --- | --- |
| pH<sup>a</sup> | 4-10 | 6-7 |
| [NaCl] (g/l)<sup>b</sup> | 0-100 | 0-10 |
| NaHCO<sub>3</sub> (mM)<sup>c</sup> | 0-300 | 0-100 |
| Temperature (°C)<sup>d</sup> | 23-45 | 37 |
| Ranges and durations: |  |  |
| <sup>a</sup>pH: 2, 4, 6, 7, 8, 10, 12 over 123 h |  |  |
| <sup>b</sup>Salinity: 0, 5, 10, 50, 80, 100 g/l over 36 h |  |  |
| <sup>c</sup>Bicarbonate: 0, 100, 300, 500 mM over 36 h |  |  |
| <sup>d</sup>Temperature: 9, 23, 27, 30, 37, 45, 55°C over 73 h |  |  |
| Growth defined as reaching OD<sub>600</sub> > 0.1 as measured by plate reader) |  |  |

#### Evaluation of growth media and substrate utilization under aerobic and 1 atm CO_2_ conditions

During growth in LB medium under 1 atm CO_2_, SR7 displayed a short lag period followed by exponential growth with a doubling time of 2.07 ± 0.1 hours until ~14 hours, when it entered stationary phase (Figure 4A). Supplementation of LB medium with 4 g/l glucose severely diminished growth (Figure 4B), possibly due to unbalanced metabolic flux through glycolysis and/or the build up of toxic metabolites resulting in premature cell death.

**Figure 4.**
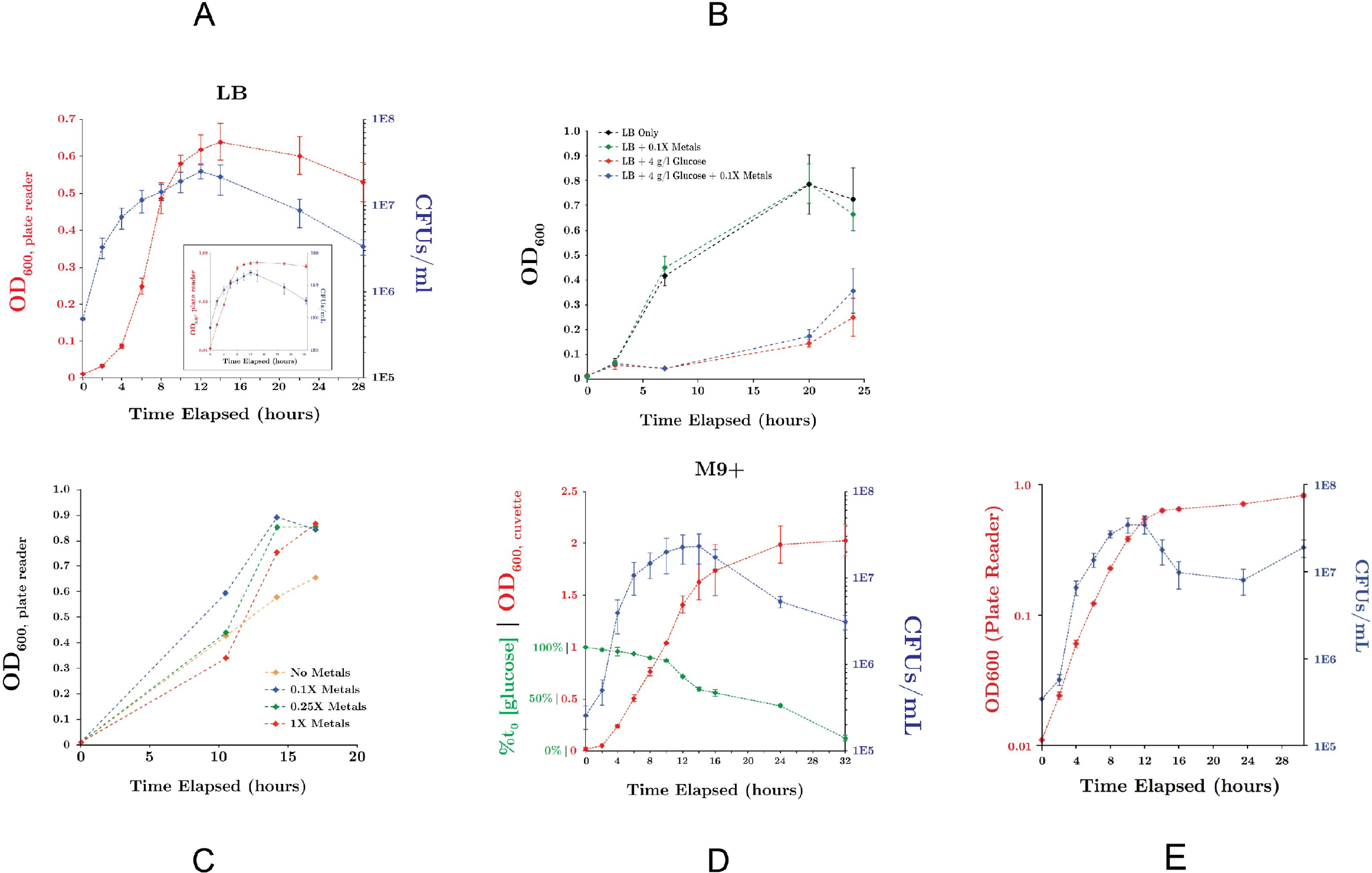
SR7 growth under 1 atm CO_2_, 37°C, 250 rpm (OD_600_: red; CFU/ml: blue; inset showing OD on logarithmic scale) **A)** in LB medium, **B)** in LB medium with glucose-amended, **C)** in M9 salts + 50 mg/l YE + 4 g/l glucose as a function of trace metals concentration, and **D)** in M9+ minimal medium (% glucose consumption: green), and **E)** in M9+ minimal media showing OD on logarithmic scale). Variation in OD_600_ ranges in plots A-C vs. D is due to different instruments used to measure optical densities (200 μl well microplate reader vs. 1 ml cuvette nanophotometer, respectively). Error bars represent standard deviation of triplicate cultures.

A minimal growth medium was developed to enable examination of microbial physiology, and nutritional growth requirements. Modification of M9 base medium with a range of carbon sources, trace metals concentrations (Figure 4C) and supplementation with dilute yeast extract (YE) enabled growth rates comparable to those observed in LB medium under 1 atm CO_2_ (Table 4). Dilute YE (50 mg/l) was insufficient to independently support observable growth, and is a common supplement to defined mineral/nutrient solutions, e.g. A5 medium (Jordan et al., 2007). Nitrate was assessed as a potential electron acceptor for anaerobic growth under 1 atm CO_2_, but did not demonstrate any pronounced effects on growth rates or biomass accumulation (Table 4).

**Table 4.**
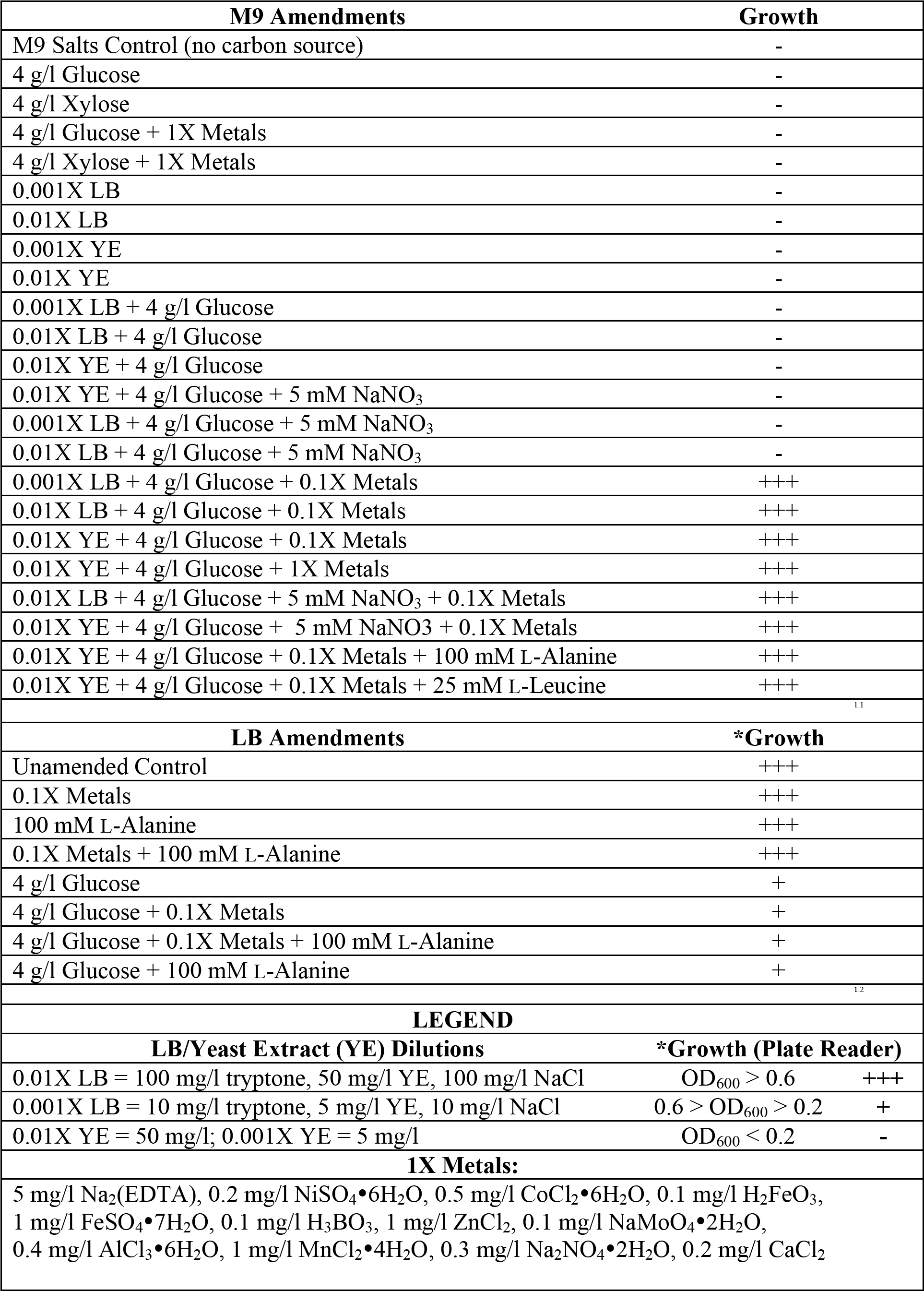
M9 and LB supplemented growth outcomes under 1 atm CO_2_

The final combination of M9 base with 50 mg/l YE, 0.1X trace metals solution and 4 g/l glucose is designated “M9+” medium, and was used as a semi-defined minimal culturing medium. SR7 displayed a doubling time of 1.93 ± 0.1 hours in M9+ medium under 1 atm CO_2_, with cultures reaching 2.0×10^7^ CFUs/ml at 12 hours (Figures 4D-E). End-product toxicity in both M9+ and LB medium may be responsible for triggering the observed decrease in viability beyond 16 hours, rather than substrate limitation. Continuous glucose consumption observed throughout stationary phase suggests that a substantial portion of the population remains metabolically active despite decreasing viable counts (Figures 4D-E). In M9+ medium, 44% and 88% of supplied glucose was consumed at 16 and 32 hours post-passaging, indicating that the carbon source was not limiting, although scarcity of other resources such as undefined YE components or trace metals may play a role in diminished population viability during late stationary phase.

Comparison by scanning electron microscopy (SEM) of mid-log phase SR7 cells grown aerobically reveals longer cells in LB medium (Figure 1D) than in M9+ (Figure 1E). Chains of vegetative cells can be observed with incomplete septal ring closures, and in some cases vegetative cells are greater than 10 μm in length (e.g. Figure 1D). Mid-log phase cells in M9+ medium under 1 atm CO_2_ (Figure 1F) appear morphologically similar to cells from aerobic conditions (Figure 1E).

The diversity of carbon substrates supporting SR7 growth was compared to two type strains of *B. megaterium*, DSM319 and QM B1551 (Tables 6 and 7). SR7 showed strong metabolic activity in minimal medium on two of 71 tested substrates: TCA Cycle intermediates citric acid and L-malic acid, while *B. megaterium* QM B1551 and DSM319 demonstrated high activity on 6 and 14 substrates, respectively (Table 7), with all three *B. megaterium* strains showing high activity on citric acid and L-malic acid (Table 7). Trace metal supplementation significantly increased the diversity of carbon substrates supporting SR7 metabolic activity from 2 to 12 (Table 6), and corroborates the positive effect of trace metal supplementation on SR7 growth rate (Figure 4). Dependence on trace metals for growth suggests that enzymes central to SR7 metabolism may require elevated metals concentrations to properly function, including calcium, manganese, cobalt and especially magnesium for biomass formation and product generation (David et al., 2010; Korneli et al., 2013a).

### Evaluation of spore germination inducers for SR7 at 1 atm CO_2_

We have previously suggested that growth of *Bacillus* spp. under scCO_2_ may be separated into two phases: 1) spore acclimation and germination, followed by 2) outgrowth of vegetative cells (Peet et al., 2015). Since fluorescence microscopy revealed that regardless of growth outcome, many ungerminated SR7 spores remain after scCO_2_ incubation, we hypothesized that increasing the likelihood and/or rapidity of SR7 spore germination would lead to observed increases in cell growth frequency under scCO_2_. Although L-alanine may appear to have moderately shortened the time to outgrowth, many SR7 spores did not germinate, as observed by CFU counts after heat kill (Figure 5A). This suggests that a subpopulation of spores germinated and grew in liquid medium, while remaining spores retain the capacity to germinate and grow, but not within the 9-hour incubation time tested. Heat activation treatment reduced SR7 germination rates and slowed doubling times (Figure 5A), despite contrary results from previous *Bacillus* spore studies (Hyatt and Levinson, 1962).

**Figure 5.**
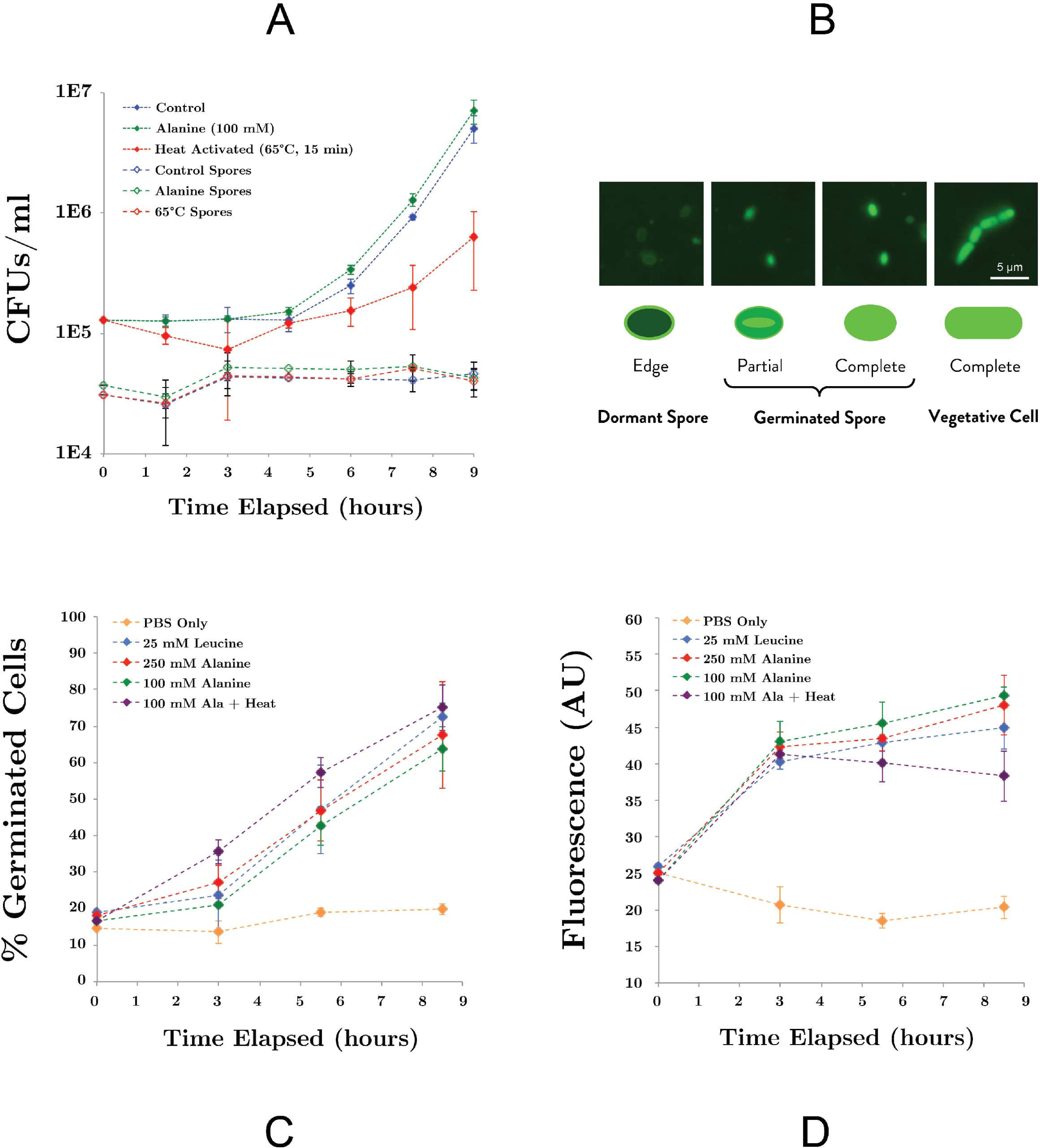
**A)** Germination of SR7 spores under 1 atm CO_2_ in LB medium, 100 mM L-alanine-amended LB, or LB heated at 65°C for 15 minutes. Open triangles representing viable spore counts based on heat-killed (80°C for 15 minutes) CFU counts. **B)** Syto9 cell staining patterns of SR7 spores and vegetative cells. ‘Edge,’ ‘Partial’ and ‘Complete’ refer to syto9 stain localization/distribution within the cell. **C)** Extent of population-wide germination in PBS incubations amended with L-alanine, L-leucine or heat activated, as measured by percent of population displaying dormant or germinated cell staining patterns assessed by microscopy. **D)** Bulk fluorescence of samples with different germination treatments.

To quantify the effect of inducers on germination rates (independent of vegetative outgrowth), SR7 spore germination was induced in PBS without any carbon source present. Fluorescence microscopy staining patterns (Figure 5B) revealed that unamended PBS-incubated spores maintained a constant, low-level abundance of germinated cells (<20% of the population), while L-leucine and L-alanine amended cultures showed a substantial increase in germination as early as 3 hours, reaching up to 75% of the population by 8.5 hours (Figure 5C). Heat treatment combined with L-alanine did not substantially increase the rate of germination, nor did increasing the L-alanine concentration from 100 mM to 250 mM. Microscopic inspection of inducer-amended PBS incubations did not reveal vegetative cell morphologies, indicating that L-alanine and L-leucine were not utilized as carbon sources for growth over the time period of this experiment. Bulk fluorescence of PBS-incubated spore suspensions generated results consistent with staining patterns; spores treated with L-alanine and L-leucine showed an approximate doubling in bulk fluorescence, while bulk fluorescence of unamended PBS cultures remained constant (Figure 5D). Since heat treatment and increased L-alanine concentrations did not enhance germination frequency, 100 mM L-alanine without heat treatment was used for the remainder of this work.

### Enhancement of SR7 growth under supercritical CO_2_ using germination inducers

Germination inducers were investigated under scCO_2_ in M9+ and phosphate buffered LB media (P-LB) using the shaking regime developed under 1 atm CO_2_ for their ability to improve SR7 vegetative growth outcomes targeting a <3 week incubation period. SR7 growth after 18-20 days under scCO_2_ was observed in 36% of unamended P-LB reactors, with all cultures generating less than 10^6^ cells/ml (Figure 6A; Table S2). Addition of L-alanine to P-LB (P-LBA) generated a maximum concentration of 2.5×10^7^ cells/ml with 63% of vessels showing growth (Figure 6A), representing a significant growth improvement over unamended P-LB cultures (Wilcoxon/Kruskal-Wallis p = 0.0036). L-alanine-amended M9+ medium (M9A+) (Figures 1G-H) supported a nearly identical growth frequency (64%) to that of P-LBA (Figure 6A), but with improved frequency of high-density (>40 fold) growth (56% in M9A+, 26% in P-LBA, 0% in P-LB; Figure 6A). Median fold increases in final cell densities relative to starting spore concentrations were 64.3, 37.5 and 22.8 for M9A+, P-LBA and P-LB cultures, respectively. In M9A+ medium, SR7 growth frequencies and fold-increases in final cell densities thus increased by factors of 1.7 (36% to 64%) and 2.8 (22.8-fold to 64.3-fold), respectively, relative to SR7 incubations in P-LB medium.

**Figure 6.**
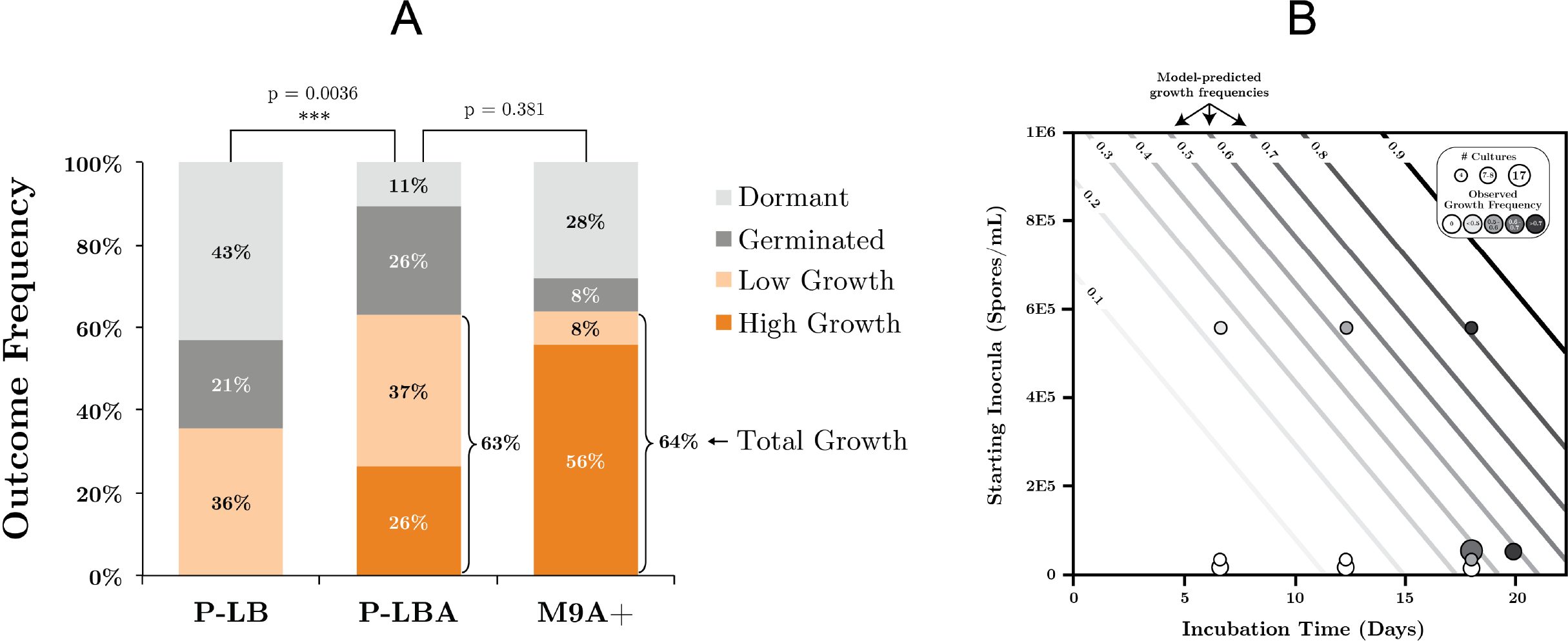
**A)** SR7 germination and growth frequencies under scCO_2_ in phosphate-buffered LB (P-LB), phosphate-buffered LB amended with l-alanine (PLBA), and M9+ amended with l-alanine (M9+A) as measured by cell quantification for multiple individual experiments. **High growth**: ≥40− fold increase in cell counts relative to t_0_; **low growth**: >10-fold increase in cell counts relative to t_0_; **germinated**: <10-fold increase, mix of vegetative cells and spores; **dormant**: <10-fold increase, spores only. All cell-free control incubations (not plotted) resulted in no growth. **B)** Logistic regression model of SR7 scCO_2_ growth outcome as a function of inocula concentration and incubation time, with predicted growth frequencies represented by contour lines (e.g. 0.5 = 50% likelihood of growth). Empirical data plotted on top of contours indicate the number of replicate cultures (symbol size) and frequency of growth (shading) for each end-point sampled scCO_2_ reactor. Growth outcomes demonstrate a statistically significant increase in the frequency of observed growth as a function of inocula spore density (p = 0.0057; Y axis) and incubation time (p = 0.003; X axis).

In contrast, L-leucine-amended cultures (P-LBL) displayed growth in only 7.7% of reactors, and the combined treatment of L-alanine and L-leucine resulted in 31% growth frequency (Table S2), suggesting that L-leucine confers a neutral to inhibitory effect on SR7 germination and growth under scCO_2_. As heat-treatment also appeared to diminish growth frequencies relative to untreated L-alanine-amended cultures (50% growth; Table S2), L-leucine and heat treatment were discarded as potential SR7 growth enhancers under scCO_2_. Due to elevated growth frequency, improved biomass accumulation, and the ability to grow from a sole carbon source under scCO_2_, M9A+ media was thus used for subsequent characterization of SR7 growth under scCO_2_. SEM analysis revealed that SR7 cells grown in M9A+ medium under scCO_2_ for 21 days (Figures 1G-H) were similar in morphology to cells grown to mid-log phase in 1 atm CO_2_ (Figure 1F) with slightly reduced cell length.

We next investigated the relationship between SR7 spore concentration and incubation time on the frequency of reactors showing positive growth, as previously conducted for *Bacillus* strains isolated from other sites (Peet et al., 2015). Logistic regression analysis (Figure 6B) established that both loaded spore density (p = 0.0057) and incubation duration (p = 0.003) have statistically significant impacts on growth frequency in M9A+ medium, while the interaction of the two variables was not significant (p = 0.89). Based on our model, in order to attain 80% growth frequency (T_80_) in vessels under scCO_2_ for SR7 in M9A+ minimal media, a starting spore concentration of 5×10^5^ spores/ml should be incubated for a minimum of 18 days (Figure 6B). This finding result represents a significant improvement over previous work where the T_80_ corresponds to 38.5 days and 37.7 days for *Bacillus* strains MIT0214 and MITOT1, respectively (Peet et al., 2015).

### Genomic characterization of isolate SR7

PacBio sequencing and assembly of *B. megaterium* SR7 DNA generated six contigs, the largest of which is 5,449,642 bp with 40.7X sequencing coverage and 39% GC content, while the other five contigs are between 2.9 kb and 22.0 kb (Table 8) and predicted to be plasmids based on gene content. Adjustments of the largest SR7 contig (Figure 7A) based on nearly 1:1 synteny (Figure 7B) and 95.9% shared average nucleotide identity (ANI) with the chromosome of reference *B. megaterium* strain QM B1551 (Eppinger et al., 2011) enabled the SR7 chromosome to be closed (Figure 7C). The six contigs of the SR7 genome are deposited under GenBank accession numbers CP022674-CP022679. Although the SR7 genome has highest ANI (97.5%) with a genome assigned to the species *Bacillus aryabhattai* (GCA_000956595), several chromosomal inversions distinguish the *B. aryabhattai* genome from *B. megaterium* SR7 (Figure 7B). As a result, SR7 has been tentatively classified as *B. megaterium* due to the high ANI (>95%) and near perfect genome synteny with other *B. megaterium* strains.

**Figure 7.**
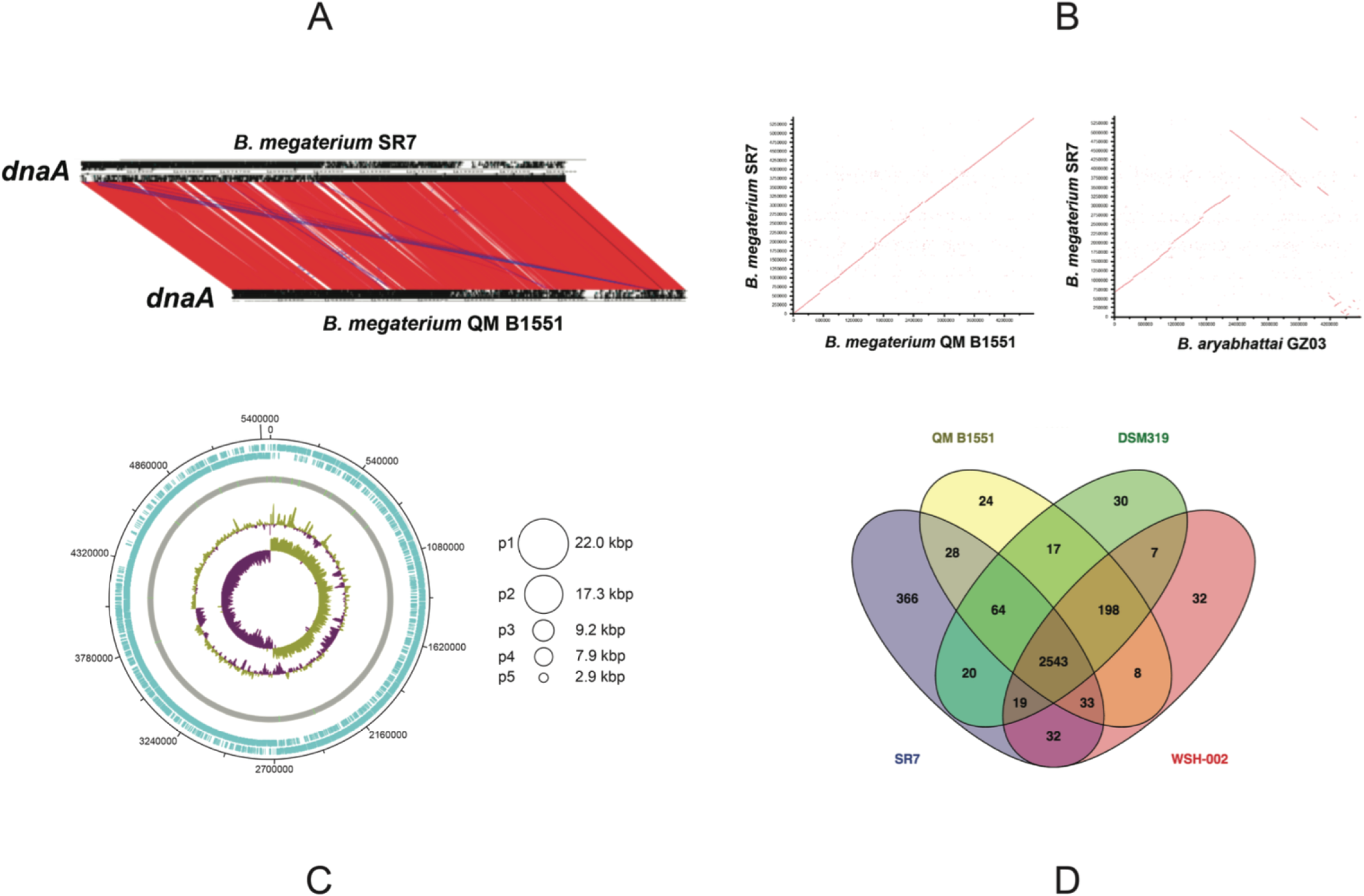
**A)** Sequence-based comparison of the largest *B. megaterium* SR7 contig to reference genome *B. megaterium* QM B1551 **B)** ANI-based comparison showing synteny between SR7 and *B. megaterium* QM B1551 and *B. aryabhattai* GZ03 **C)** Schematic of the *B. megaterium SR7* 5.51 Mbp genome, including the closed 5.45 Mbp chromosome. Concentric circles (outside in) are RAST ORFs (blue), rRNA and tRNA (green lines on grey circle), GC content, and GC skew. Asymmetry in GC skew indicates proper chromosome assembly (Grigoriev, 1998). Circles at right represent five putative plasmids (p1-p5) native to SR7. **D)** Venn diagram displaying shared RAST-called and annotated ORFs between SR7 and three previously sequenced *B. megaterium* strains. 366 RAST-called ORF annotations unique to SR7 are listed in Table S3.

Gene prediction via RAST (Overbeek et al., 2014) indicated 5,696 genes, with 13 complete rRNA operons with 5S, 16S and 23S rRNA genes. The *B. megaterium* SR7 genome size (5.51 Mbp) is 7.5% larger than that of QM B1551 (5.1 Mbp) and DSM319 (5.1 Mbp) (Eppinger et al., 2011), and approximately 33% larger than strain WSH-002 (4.14 Mbp) (Liu et al., 2011). A RAST-based comparison of gene content revealed that 2,543 annotated genes (82% of annotated SR7 ORFs) are shared by all four sequenced *B. megaterium* genomes (SR7, DSM319, QM B1551, WSH-002) (Figure 7D) while SR7 has 366 gene annotations (12% of SR7 ORFs) that are unique to the strain (Table S3). 59.2 kbp (1.1%) of SR7’s genomic content is located on its five putative plasmids, which is consistent with previous studies establishing the propensity for *B. megaterium* strains to store large extra chromosomal DNA in plasmids (Eppinger et al., 2011; Liu et al., 2011; Korneli et al., 2013b). Several regions of synteny were observed between the five smaller SR7 plasmid-like contigs and QM B1551 plasmids; most of the ORFs from these small SR7 contigs were annotated as hypothetical proteins or displayed highest homology to genes associated with plasmid replication, recombination, sporulation, and motility, and/or plasmid-borne genes from *Bacillus spp*. including *B. megaterium* (Table S4), leading to our classifying them as plasmids p1 through p5 (Figure 7C).

Functional annotation of the SR7 genome (Table 5) includes genes associated with complete glycolysis, Entner-Doudoroff Pathway, TCA Cycle, Pentose Phosphate Pathway, Glyoxylate Bypass, and biosynthesis pathways for all amino acids. Chromosomal annotation also includes broad capacity for sugar transport and utilization, including for glucose, fructose, mannose, and xylose, and the production of butyrate, lactate, butanol, acetate, 2,3-butanediol, succinate and ethanol. Annotations associated with anaerobic respiration and redox metabolism include dissimilatory sulfate (*sat*), nitrite reduction (*nirBD*) and denitrification (*norQD*). No genes associated with carbon fixation pathways (i.e. Calvin Cycle, Wood-Ljungdahl Pathway, rTCA cycle, etc.) were detected in the genome. However, the annotation of carbonic anhydrase (converts CO_2_ to H_2_CO_3_) (Smith and Ferry, 2000), carbamoyl-phosphate synthase (incorporates bicarbonate for pyrimidine and arginine biosynthesis) (Arioli et al., 2009), and several carboxylases including phosphoenolpyruvate carboxylase (catalyzes the irreversible addition of bicarbonate to phosphoenolpyruvate) (Din et al., 1967; Santillan et al., 2015) indicates the capacity for SR7 to assimilate gaseous and dissolved CO_2_ species. The presence of carboxylases may prove useful for future engineering of CO_2_-consuming metabolic pathways under scCO_2_ headspace, especially in light of the previous demonstration of *B. megaterium* carboxylase activity under scCO_2_ (Matsuda et al., 2001).

**Table 5.**
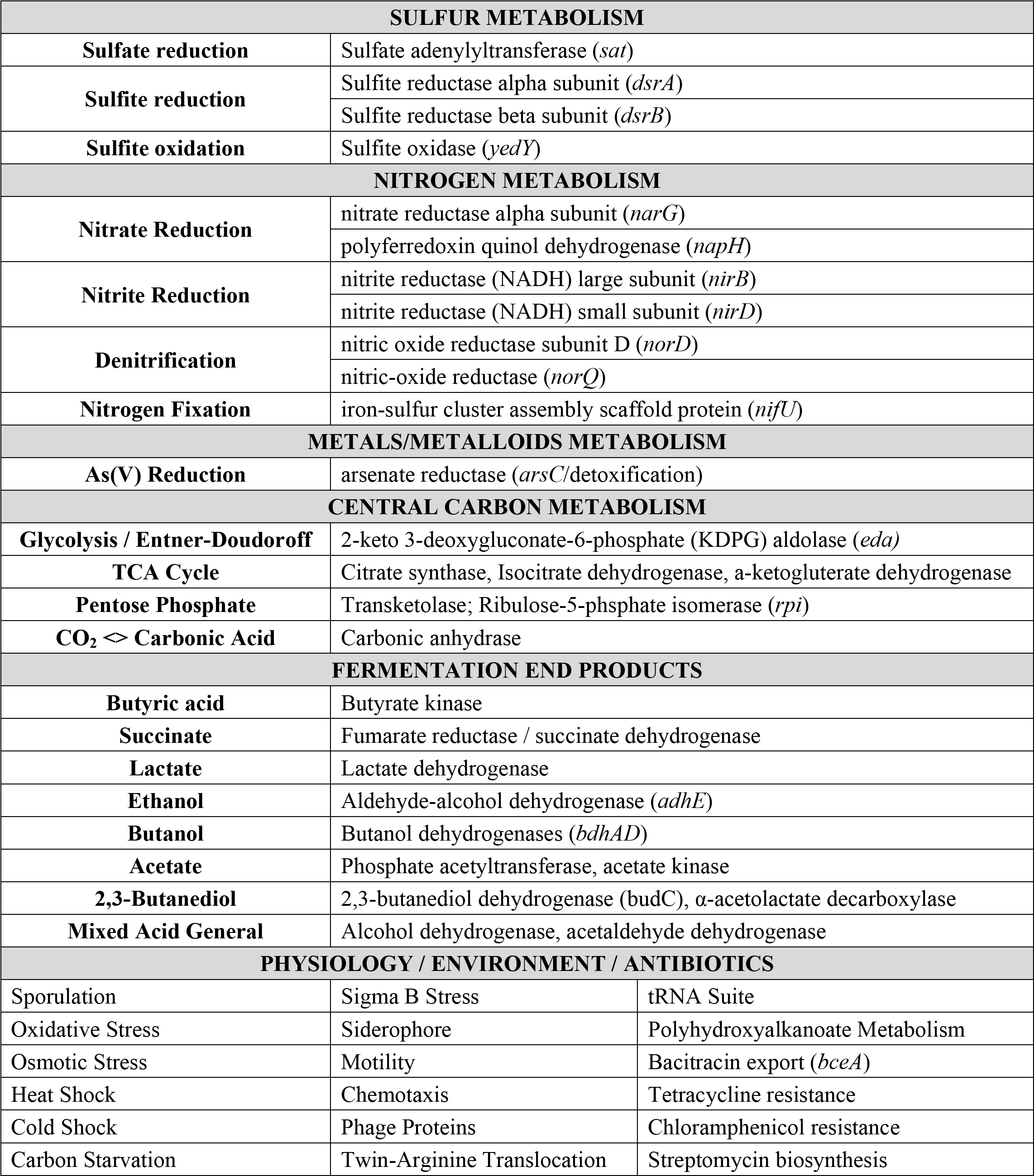
Summary of isolate *B. megaterium* SR7 functional genomic annotation (RAST)

Several additional genes and pathways predicted from the SR7 genome are worth noting for their potential utility as components of a microbial bioproduction system, including sporulation, siderophore assembly and uptake, flagellar motility, the twin-arginine translocation (TAT) system, and PHA synthesis, the last of which indicates a capacity for redirecting flux toward concentrated storage of excess carbon (Table 5). The endogenous TAT secretion system may be useful for developing the ability to secrete proteins or enzymes as engineered bioproducts into aqueous or scCO_2_ phases, including those capable of cell signaling or degrading lignocellulosic feedstocks (Kikuchi et al., 2008; Schaerlaekens et al., 2004; Anné et al., 2012). In addition, the SR7 chromosome and putative plasmids were searched using the virulence factor database (Chen et al., 2016). No toxins were annotated on any of the SR7 plasmids or genome, and all proteins homologous to known virulence factors are also associated with functions unrelated to virulence (e.g. *capABCD* for capsid synthesis, flagellar genes, and clp stress-response proteins).

### Characterization of SR7 fermentation products under 1 atm CO_2_ and supercritical CO_2_

Several fermentation pathways were predicted in the annotated SR7 genome and validated by detection of their end products in SR7 cultures. Genes for L-lactate dehydrogenase (*ldhA*) for lactate formation; phosphate acetyltransferase (*pck*), acetate kinase (*ackA*) and acetyl-CoA synthase (*acs*) for acetate production and reassimilation; eight alcohol dehydrogenases; and a complete set of TCA Cycle enzymes (including succinyl-CoA synthetase), indicate the capacity for SR7 to generate ethanol and succinate from glucose (Table 5). Mid-log phase cultures of SR7 passaged under 1 atm CO_2_ generated acetate in both LB and M9+ medium (after 6 hours: 271 mg/l in LB; 49 mg/l in M9+; Figures 8A-B), while LB cultures also generated succinate (300 mg/l after 6 hours). In M9+ medium, stationary phase cultures at 1 atm CO_2_ (48 hours after passaging) continue to generate acetate (352 mg/l), with large amounts of lactate (1.54 g/l), and low concentrations of succinate (62 mg/l) and ethanol (17 mg/l) (Figure 8B). In LB medium, stationary-phase cultures generated a metabolite profile constrained to succinate (1.11 g/l) and acetate (235 mg/l; Figure 8A). Fermentative compounds produced by SR7 under 1 atm CO_2_ in M9+ medium imply glycolytic conversion of glucose to pyruvate, followed by production of lactate, acetate, succinate, or ethanol. The generation of succinate and acetate in LB medium indicates expression of pathways catabolizing undefined yeast extract components that funnel into similar fermentative steps.

**Figure 8.**
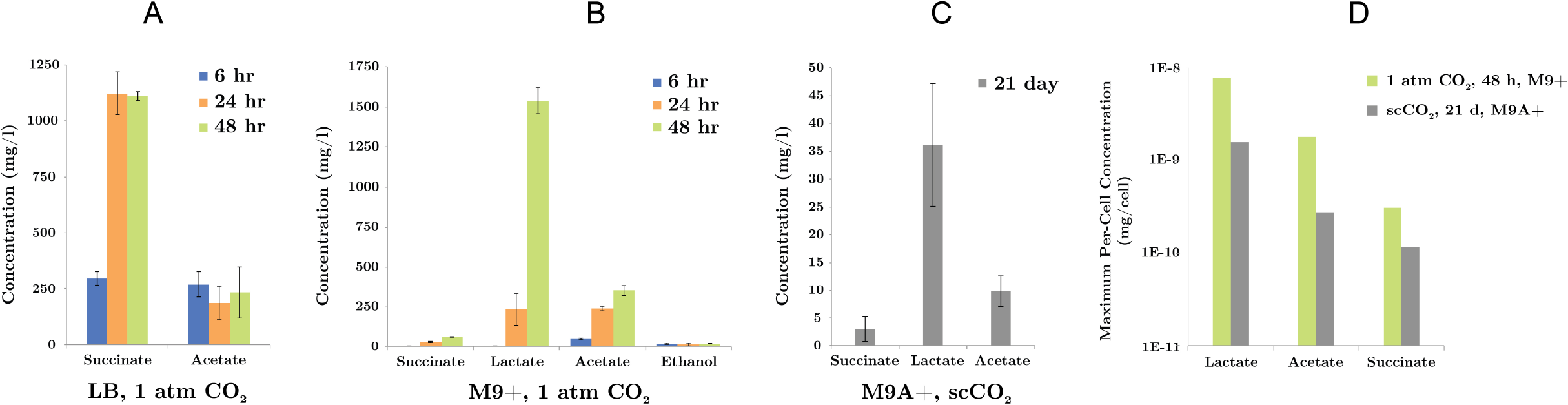
SR7 fermentation products (mg/l) under 1 atm CO_2_ in **A)** LB medium and **B)** M9+ medium (sampled 6, 24, and 48 hours after passaging of vegetative cultures) and **C)** under scCO_2_ in M9A+ medium for spore-inoculated cultures demonstrating growth (end-point sampled at 21 days). **D)** Maximum per-cell fermentation product concentrations (mg/l) for SR7 cultures demonstrating growth under 1 atm CO_2_ and scCO_2_ in M9+ and M9A+ media, respectively. Per-cell normalization based on cell densities enumerated by fluorescence microscopy. No fermentation products were detected in cell-free control incubations or scCO_2_ cultures not demonstrating growth. Error bars represent standard deviation of triplicate cultures.

Fermentation products generated in scCO_2_ cultures grown in M9A+ medium, including lactate (up to 47.6 mg/l), acetate (up to 13.2 mg/l), and succinate (up to 5.4 mg/l) (Figure 8C), were similar to those observed in M9+ under 1 atm CO_2_, suggesting similar metabolic activity under both conditions. Importantly, detection of fermentation products represents the first evidence of bioproduct generation by actively growing cultures under scCO_2_. Maximum fermentation product concentrations generated in M9A+ medium after 21 days under scCO_2_ are lower than for 1 atm cultures that have reached late stationary phase at 48 hours. However, maximum per-cell titers under scCO_2_ in M9A+ medium of 1.6×10^−9^, 2.8×10^−10^, and 1.2×10^−10^ mg/l for lactate, acetate, and succinate, respectively, are comparable to per-cell results under 1 atm CO_2_ in M9+ media, i.e. 7.7×10^−9^, 1.8×10^−9^ and 3.1×10^−10^ mg/cell for lactate, acetate, and succinate, respectively (Figure 8D). Maximum per-cell concentrations are thus within roughly one order of magnitude for SR7 cultures incubated under 1 atm CO_2_ and scCO_2_ (with alanine supplementation), with higher fermentative output observed for cultures incubated under 1 atm CO_2_ conditions.

## Discussion

Supercritical CO_2_ is widely used as an extractive solvent in chemical industrial processes. Its potential utility for bioproduct harvesting has been recognized, but not realized due to a lack of compatibility with established systems for microbial bioproduction. The development of a metabolically active organism under scCO_2_ with broad engineering potential would thus represent a major step toward the direct use of scCO_2_ in bioprocessing, including as a sterilizing phase or as a solvent for *in situ* bioproduct extraction. Recently, we have shown that multiple *Bacillus* strains are capable of growth under scCO_2_ following inoculation as spores (Peet et al, 2015), opening the possibility of using *Bacillus spp*. as biocatalysis or bioproduction agents coupled with *in situ* extraction by scCO_2_. In this work we have employed a bioprospecting approach to identify and isolate a scCO_2_-biocompatible *B. megaterium* strain that demonstrates higher growth frequencies after shorter incubation periods than previously isolated strains. Through minimal medium development, culturing optimization and induction of germination with L-alanine, we improved growth probabilities for <3 week incubations by 1.7-fold, increased final cell densities relative to starting spore concentrations by 2.8-fold, and demonstrated, for the first time, fermentation product generation under scCO_2_. The fermentation products under both 1 atm CO_2_ and scCO_2_ were consistent with annotated metabolic pathways in the SR7 genome and provide proof of concept that *in vivo* generation of extracellular compounds is possible in aqueous phase cultures under scCO_2_.

In steel columns, microbial growth from spores under scCO_2_ is thought to occur in two distinct phases (Peet et al., 2015). First, dormant scCO_2_-resistant spores may become acclimated to the high pressure and acidity (due to elevated levels of carbonic acid). Second, a subpopulation of acclimated spores may germinate stochastically and commence outgrowth as vegetative cells, allowing a subset of spores to initiate metabolic activity in response to changing or stressful environmental conditions without putting the remaining population at risk of sterilization (Buerger et al., 2012). Therefore, in addition to having bioprospected SR7 for its native resilience to SR7 exposure, development of culture conditions that support *B. megaterium* SR7 growth targeted two key steps – spore germination and vegetative outgrowth – in achieving enhanced growth frequencies and biomass accumulation under scCO_2_. SR7 germination rates and doubling times were initially improved under 1 atm CO_2_ by increased mixing speeds (Figure 3), which follows trends in the literature showing a correlation between *B. megaterium* growth rates and disruptive mixing (Santos et al., 2014). While this correlation has previously been linked to media aeration, here we show mixing rate influences growth under anaerobic conditions as well, potentially due to mass transfer of inhibitor end products or surface attachment effects.

Literature on *Bacillus* physiology (Roth and Lively, 1956; Hyatt and Levinson, 1962; Levinson and Hyatt, 1970; Vary, 1973; Cronin and Wilkinson, 2007; Wei et al., 2010; Setlow, 2014) elucidated precedent for inducing spore germination through conditional (e.g. pressure; temperature; (Wei et al., 2010)) or chemical (Ghosh and Setlow, 2009) treatments. We thus hypothesized that amending growth medium with a germination inducer or subjecting cultures to heat activation would facilitate more rapid and consistent vegetative outgrowth under scCO_2_ relative to untreated cultures subject to diminished rates of stochastic germination. Our observations were consistent with this hypothesis for the inducer L-alanine, but not for L-leucine or heat treatment.

Initial culturing of SR7 spores under 1 atm CO_2_ suggested that addition of the germination inducer L-alanine hastened vegetative outgrowth (Figure 5A) and initiated cellular modifications consistent with germination (Figures 5B-D). Results from sublethal heat activation treatment experiments were partially consistent with chemical inducers, though may reduce growth viability in sub-populations, possibly through spore-coat damage (Figure 5A) (Setlow, 2006). Regardless of treatment, spore concentrations under 1 atm CO_2_ remained nearly constant (Figure 5A) despite initiation of vegetative outgrowth, reinforcing the notion that under a CO_2_ atmosphere, spore subpopulations trigger active metabolism, while the majority of spores remain dormant. While addition of L-alanine may therefore increase the potential for a pool of dormant cells to germinate, initial results under 1 atm CO_2_ were less dramatic than results under high pressure scCO_2_ conditions.

Addition of L-alanine under scCO_2_ enabled a statistically significant improvement in high-pressure growth outcomes in both P-LBA and M9A+ medium compared to P-LB (Figure 6A). Relative to *Bacillus* strains MIT0214 and MITOT1, which displayed 33% to 55% scCO_2_ growth frequencies, respectively, after ≥20 days in complex medium when inoculated with 10^4^ spores/ml (Peet et al., 2015), SR7 outperformed these strains using the same inoculum concentration by demonstrating 64% growth frequency after 18-20 days in M9A+. The generation of a nominal logistic regression model (Figure 6B) provides mathematical predictive power regarding growth outcomes (i.e. inoculation with 5×10^5^ spores/ml is expected to demonstrate 80% growth frequency over 18 days), which informs the design and duration of future scCO_2_ culturing experiments and provides a benchmark against which to measure system improvements. Previous studies (Vary, 1973; Wei et al., 2010) have shown that the likelihood of germination may be a function of spore age or repeated cycles of germination and sporulation, with an undesired state of “superdormancy” resulting in diminished likelihood of vegetative outgrowth. Since the initiation of germination may also increase as a function of spore age in the presence of different chemical inducers and buffer concentrations (Vary, 1973), a combinatorial study of spore storage duration with a range of alternative inducers and culturing media may be of particular utility. It will also be helpful in future work to explore whether sub-populations of spores respond differently to scCO_2_ during germination (with or without induction) with the ultimate goal of creating homogenous spore populations for more consistent outgrowth.

While low frequency growth (Peet et al., 2015) and enzymatic biocatalysis (Matsuda et al., 2001) have previously been shown under scCO_2_, this work describes the first combined demonstration of enhanced high-density vegetative growth and endogenous metabolite production under scCO_2_ conditions (Figure 8C). While acetate, lactate and succinate do not partition effectively into scCO_2_, their detection demonstrates that extracellular compound production is feasible for SR7, possibly enabling *in situ* extraction for compounds that are preferentially extracted by scCO_2_ from the aqueous phase. Though strain SR7 generated only trace amounts of ethanol in M9+ medium under 1 atm CO_2_, in the event of higher titers under scCO_2_ in alternative medium or by a different host organism (e.g. *Clostridium thermocellum*; Knutson et al., 1999), the effective partitioning of ethanol into the scCO_2_ phase (Guvenc et al., 1998) would prolong batch culture growth capacity by reducing the aqueous concentration of this common toxic end-product (Herrero, 1983).

Isolation of SR7 is especially fortunate due to the extensive research performed on *B. megaterium* as a bioproduction host. Three *B. megaterium* strain genomes have been fully sequenced and annotated (Eppinger et al., 2011; Liu et al., 2011) in addition to a genome scale model for WSH-002 (Zou et al., 2013) that enabled comparative genomics with SR7. Numerous systems biology (Biedendieck et al., 2011), metabolic engineering and synthetic biology (Marchand and Collins, 2016; Moore et al., 2018) tools are available for type strains of *B. megaterium*, potentially extending the biotechnological utility of SR7, as well as having methods available to further study its metabolism and physiology, e.g. other strains of *B. megaterium* have recently been engineered to produce high titers of Vitamin B12 (Moore et al., 2014), shikimic acid (Ghosh and Banerjee, 2015), and levansucrase (Korneli et al., 2013a). *B. megaterium* is also known to secrete significant amounts of enzymes and proteins (Korneli et al., 2013b), enabling degradation of complex feedstock components into simpler substrates for intracellular transport and consumption (Bunk et al., 2010). Consistent with the capacity to metabolize a wide variety of compounds (Vary, 1994), strain SR7 demonstrated growth on a broad substrate spectrum, including organic acids, sugars and amino acids (Table 6). The elevated number of rRNA genes (13) appears consistent with previous findings that spore-forming soil bacteria have some of the highest observed numbers of rRNA operons per genome (Klappenbach et al., 2000).

**Table 6.** SR7 robust (+++), marginal (+), and no (−) growth on sole carbon source in unamended control and trace metals-amended Biolog plates. Maximum growth per plate noted by an asterisk

| Carbon Substrate | Control | +Metals |
| --- | --- | --- |
| Citric Acid | +++* | +++ |
| α-D-Glucose | + | +++ |
| L-Arginine | + | +++ |
| D-Gluconic Acid | + | +++ |
| L-Aspartic Acid | + | +++ |
| N-Acetyl-D- Glucosamine | + | +++* |
| L-Glutamic Acid | + | +++ |
| D-Turanose | + | +++ |
| L-Pyroglutamic Acid | + | +++ |
| D-Raffinose | + | +++ |
| γ-Amino-Butyric Acid | + | +++ |
| myo-Inositol | + | +++ |
| L-Malic Acid | +++ | + |
| Gelatin | + | + |
| Pectin | + | + |
| Dextrin | + | + |
| α-D-Lactose | + | + |
| D-Mannitol | + | + |
| Methyl Pyruvate | + | + |
| D-Melibiose | + | + |
| D-Fructose | + | + |
| D-Arabitol | + | + |
| L-Alanine | + | + |
| D-Lactic Acid Methyl Ester | + | + |
| D-Galactose | + | + |
| L-Lactic Acid | + | + |
| β-Hydroxy-D,L- Butyric Acid | + | + |
| D-Cellobiose | + | + |
| D-Salicin | + | + |
| Glycerol | + | + |
| Gentiobiose | + | + |
| α-Keto-Glutaric Acid | + | + |
| L-Histidine | + | + |
| Stachyose | + | + |
| Bromo-Succinic Acid | + | + |
| D-Maltose | + | + |
| D-Trehalose | + | + |
| β-Methyl-D-Glucoside | + | + |
| Sucrose | + | + |
| Inosine | + | + |
| D-Sorbitol | - | + |
| 3-Methyl Glucose | - | + |
| D-Glucuronic Acid | + | - |
| Acetic Acid | + | - |
| L-Serine | + | - |
| Tween 40 | + | - |
| D-Galacturonic Acid | + | - |
| L-Galactonic Acid Lactone | + | - |
| Acetoacetic Acid | + | - |
| Mucic Acid | + | - |
| Propionic Acid | + | - |
| Quinic Acid | + | - |
| D-Saccharic Acid | + | - |
| D-Fructose- 6-PO<sub>4</sub> | - | - |
| N-Acetyl-D- Galactosamine | - | - |
| Formic Acid | - | - |
| Negative Control | - | - |
| p-Hydroxy- Phenylacetic Acid | - | - |
| D-Mannose | - | - |
| Glycyl-L-Proline | - | - |
| α-Hydroxy- Butyric Acid | - | - |
| α-Keto-Butyric Acid | - | - |
| D-Fucose | - | - |
| D-Glucose- 6-PO<sub>4</sub> | - | - |
| Glucuronamide | - | - |
| N-Acetyl-β-D- Mannosamine | - | - |
| L-Fucose | - | - |
| D-Malic Acid | - | - |
| L-Rhamnose | - | - |
| D-Aspartic Acid | - | - |
| N-Acetyl Neuraminic Acid | - | - |
| D-Serine | - | - |

**Table 7.** *B. megaterium* strains categorized by robust (+++), marginal (+), or no (−) growth on Biolog sole carbon sources (no metals added). Substrates enabling robust growth by at least one strain listed.

| Carbon Substrate | SR7 | DSM319 | QM B1551 |
| --- | --- | --- | --- |
| Citric Acid | +++ | +++ | +++ |
| L-Malic Acid | +++ | +++ | +++ |
| L-Lactic Acid | + | +++ | +++ |
| L-Glutamic Acid | + | +++ | +++ |
| α-D-Glucose | + | +++ | + |
| Dextrin | + | +++ | + |
| D-Mannitol | + | +++ | + |
| D-Gluconic Acid | + | +++ | + |
| L-Aspartic Acid | + | +++ | + |
| N-Acetyl-D- Glucosamine | + | +++ | + |
| L-Histidine | + | +++ | + |
| Bromo-Succinic Acid | + | +++ | + |
| D-Maltose | - | +++ | + |
| Sucrose | - | +++ | + |
| β-Hydroxy-D,L-Butyric Acid | + | + | +++ |
| D-Saccharic Acid | - | - | +++ |

Because *Bacillus spp*. have previously been detected in natural (Mu et al., 2014; Freedman et al., 2017) and laboratory (Peet et al., 2015) biphasic aqueous-scCO_2_ systems, it is unsurprising that serial passaging of deep subsurface McElmo Dome scCO_2_ reservoir formation fluids enabled the isolation of another spore-forming *Bacillus* strain in SR7. The isolation of an aerotolerant, spore-forming strain also appears reasonable due to stresses associated with sampling, including depressurization, brief oxygen exposure, and chilled temperatures during transport and storage. Physiological characterization of SR7 established viable growth ranges for pH and salinity (Table 3) consistent with *in situ* deep subsurface conditions at McElmo Dome (Morgan et al., 2001). Detection of *Bacillus spp*. in McElmo Dome community surveys (Freedman et al., 2017) and other deep subsurface formations (Vary, 1994; Nicholson, 2002) support *in situ* habitation. However, while McElmo Dome fluid temperatures are typically >60°C, we found 45°C as an upper temperature boundary for vegetative SR7 growth (Table 3), with spores remaining viable at 65°C. These results suggest that SR7 may persist at McElmo Dome as dormant endospores with the potential to acclimate to elevated temperatures upon germination.

The specific cellular tolerance mechanisms to scCO_2_ exposure remain largely unknown, although prior work suggests modifications to proteomic signatures, membrane lipid content (Budisa and Schulze-Makuch, 2014) and extrapolymeric substance (EPS) surface layer production (Mitchell et al., 2008) may be implicated in scCO_2_ survival. In addition, resilience to initial scCO_2_ exposure has been demonstrated for some Gram-positive bacteria (e.g. *Bacillus spp.*) and biofilm-associated bacteria, where thick peptidoglycan cell walls and EPS layers may support a bacterial microenvironment protective of stresses from scCO_2_ (Dhami et al., 2013; Mitchell et al., 2008). Bacterial spores, in particular those of Bacilli, are known especially in the food industry to be resistant to scCO_2_ treatment when attached to stainless steel piping, although germination and growth is not observed (Park et al., 2013). A key difference between those experiments and the laboratory conditions described in this study is the presence of an aqueous phase, where it is believed the acclimation process that enables vegetative outgrowth is occurring. Previous work has shown that without the presence of an aqueous phase, spores are able to withstand scCO_2_ exposure to a lesser degree and are unable to grow (Peet et al., 2015). As with surface attachment behavior of the related species *B. anthracis* (Williams et al., 2013), adherence to stainless steel columns may provide SR7 stabilizing benefits that enhance acclimation capacity resulting in initiation of germination upon attachment.

**Table 8.** SR7 genome sequencing, assembly, and annotation statistics

| Contig # | DNA Type | Length (bp) | % Bases Called | Fold Coverage | <sup>1</sup>Total ORFs | <sup>2</sup>Plasmid-Associated | <sup>3</sup>Sporulation/Germination |
| --- | --- | --- | --- | --- | --- | --- | --- |
| 1 | Chromosome | 5449642 | 100.0 | 40.7 | 5,696 | 3 | 194 |
| 2 | Plasmid p1 | 21958 | 99.9 | 57.0 | 35 | 11 | 6 |
| 3 | Plasmid p2 | 17283 | 100.0 | 50.5 | 19 | 4 | 2 |
| 4 | Plasmid p3 | 9202 | 79.2 | 20.8 | 13 | 3 | 1 |
| 5 | Plasmid p4 | 7873 | 92.5 | 6.7 | 8 | 3 | 1 |
| 6 | Plasmid p5 | 2921 | 52.2 | 0.5 | 4 | 2 | 1 |

<sup>1</sup>RAST-called and annotated (http://rast.nmpdr.org/)<sup>2</sup>Blastp-annotated: plasmid-associated function or previously detected on *Bacillus spp.* plasmids<sup>3</sup>Blastp-annotated: sporulation/germination-associated function

While this study has focused on increasing the frequency of growth under scCO_2_, we continually seek methods to accelerate SR7 acclimation and outgrowth under scCO_2_, including optimized reactor design, spore inoculum preparations (Prentice et al., 1972), and fine-tuned media development. Future work will also include further definition of minimal media formulations to enable analyses of metabolic flux. Characterization of SR7 under varying CO_2_ pressures may aid in identifying exploitable traits specific to scCO_2_ tolerance. For example, introduction of the tryptophan permease gene *TAT2* into *S. cerevisiae* was shown to confer high-pressure tolerance, enabling growth of yeast under pressures reaching nearly 250 atm (Abe and Horikoshi, 2000). Improved fitness may also be achieved through strain evolution, which in the context of scCO_2_ may center on high-pressure and low pH acclimation. Similarly, *E. coli* subjected to increasingly high-pressure treatments generated a piezotolerant mutant strain capable of growth at >500 atm (Nguyen, 2013). Beyond SR7 performance, this approach could be compelling for strains already capable of generating scCO_2_-compatible compounds (e.g. through ABE fermentation), including spore-forming *Clostridia* species *C. cellulovorans* (Yang et al., 2015), *C. thermocellum* (Knutson et al., 1999) or *C. acetobutylicum* (Lee et al., 2012).

Whether it is with *B. megaterium* SR7 or an alternative species, reactor incubations using a scCO_2_-resistant bioproduction strain holds strong potential for mitigating contamination concerns due to broad microbial lethality of scCO_2_ exposure, and may relieve end-product toxicity by continuous removal of extracellular compounds that preferentially partition into the scCO_2_ phase. Moreover, *in vivo* microbial bioproduction would facilitate enzyme regeneration and protection from scCO_2_ degradation, improving the consistency of biocatalysis relative to *in vitro* reactions while reducing the cost and burden of enzyme supplementation. As short-to-medium chain alcohols readily partition into the scCO_2_ phase from aqueous solutions (Laitinen and Kaunisto, 1999; Timko et al., 2004) and have been produced extensively in *Bacillus* species (e.g. Nielsen et al., 2009), advanced biofuels represent a potentially exciting class of microbially-generated compounds within a scCO_2_ bioproduction system. CO_2_-fixing carboxylation reactions are also of interest due to their potential to reduce dependence on expensive carbohydrate-based feedstocks and precedent for *in vitro* enzymatic fixation of scCO_2_ solvent (Sugimura et al., 1989; 1990; Hartman and Harpel, 1994; Kawanami and Ikushima, 2000; Yoshida et al., 2000; Matsuda, 2013), including by utilizing purified decarboxylases from *Bacillus megaterium* (Wieser et al., 1998; Yoshida et al., 2000). With these goals in mind, attention may now turn to SR7 genetic system development, heterologous pathway engineering, and optimized bioreactor design as a means for improving existing methods of aseptic culturing, end-product inhibition relief, and biocatalyzed product generation and extraction.

## Acknowledgements

The authors wish to thank members of the Thompson and Prather labs for helpful discussions and feedback. We thank KinderMorgan for access to fluid-water separators and assistance with sampling at the McElmo Dome field, CO. We gratefully acknowledge genome sequencing support by the MIT BioMicro Center, SEM imaging assistance by Nicki Watson at the Whitehead Institute for Biomedical Research, and the Chisholm and Polz labs at MIT for access to their epifluorescence microscopy facility. This material is based upon work partially supported by the U.S. Department of Energy, Office of Science, Office of Biological and Environmental Research under Award Number DE-SC0012555. Support for AJEF was also provided by the MIT Department of Civil and Environmental Engineering and the NIH/NIGMS Interdepartmental Biotechnology Training Program (award GMS T32GM008334). Support for sequencing was provided by the MIT Center for Environmental Health Science BioMicro Center, NIEHS Grant #P30-ES002109.

## Supplementary material

Supplementary Material is available at the Frontiers in Microbiology website.

## Author Contributions

**AF** was responsible for strain isolation, characterization, and growth optimization under CO_2_ headspace, data analysis, and drafting the manuscript. **KCP** assisted with enrichment culture and passaging of environmental samples and cell quantification by fluorescence microscopy. **JB** aided in strain development under aerobic conditions, including media optimization, Biolog phenotypic fingerprinting and HPLC analysis. **KP** helped with PacBio genome sequencing and genome closure. **KLJP** and **JT** advised on design for experimental and analytical approaches and **JT** supervised the project. All authors provided critical feedback and final approval of the manuscript, and agree to be accountable for all aspects of the published work.

## Disclaimer

“This publication was prepared as an account of work sponsored by an agency of the United States Government. Neither the United States Government nor any agency thereof, nor any of their employees, makes any warranty, express or implied, or assumes any legal liability or responsibility for the accuracy, completeness, or usefulness of any information, apparatus, product, or process disclosed, or represents that its use would not infringe privately owned rights. Reference herein to any specific commercial product, process, or service by trade name, trademark, manufacturer, or otherwise does not necessarily constitute or imply its endorsement, recommendation, or favoring by the United States Government or any agency thereof. The views and opinions of authors expressed herein do not necessarily state or reflect those of the United States Government or any agency thereof.”

